# Pharmacological depletion of RNA splicing factor RBM39 by indisulam synergizes with PARP inhibitors in high-grade serous ovarian carcinoma

**DOI:** 10.1101/2023.01.18.524417

**Authors:** Yuewei Xu, Sarah Spear, Yurui Ma, Marc P. Lorentzen, Michael Gruet, Flora McKinney, Yitao Xu, Chiharu Wickremesinghe, Madelen R Shepherd, Iain McNeish, Hector C. Keun, Anke Nijhuis

**Affiliations:** Department of Surgery & Cancer, Imperial College London; London, UK; Ovarian Cancer Action Research Centre, Department of Surgery & Cancer, Imperial College London, London UK

## Abstract

Ovarian high-grade serous carcinoma (HGSC) is the most common and lethal subtype of ovarian cancer with limited therapeutic options. In recent years, PARP inhibitors have demonstrated significant clinical benefits, especially in patients with BRCA1/2 mutations. However, acquired drug resistance and relapse is a major challenge. Therapies disrupting the spliceosome alter cancer transcriptomes and have shown potential to improve PARP inhibitor response. Indisulam (E7070) has been identified as a molecular glue that brings splicing factor RBM39 and DCAF15 E3 ubiquitin ligase in close proximity. Exposure to indisulam induces RBM39 proteasomal degradation through DCAF15-mediated polyubiquitination and subsequent RNA splicing defects. In this study, we demonstrate that loss of RBM39 induces splicing errors in DNA damage repair genes in ovarian cancer, leading to increased sensitivity to PARP inhibitors such as olaparib. Indisulam synergized with olaparib in multiple *in vitro* models of ovarian cancer regardless of PARP inhibitor sensitivity and improved olaparib response in mice bearing PARP inhibitor-resistant tumors. DCAF15 expression, but not *BRCA1/2* mutational status, was essential for the synergy between indisulam and olaparib, suggesting that the combination therapy may benefit patients irrespective of their *BRCA1/2* status. These findings demonstrate that combining RBM39 degraders and PARP inhibitors is a promising therapeutic approach to improving PARP inhibitor response in ovarian HGSC.

**Graphical Abstract:** 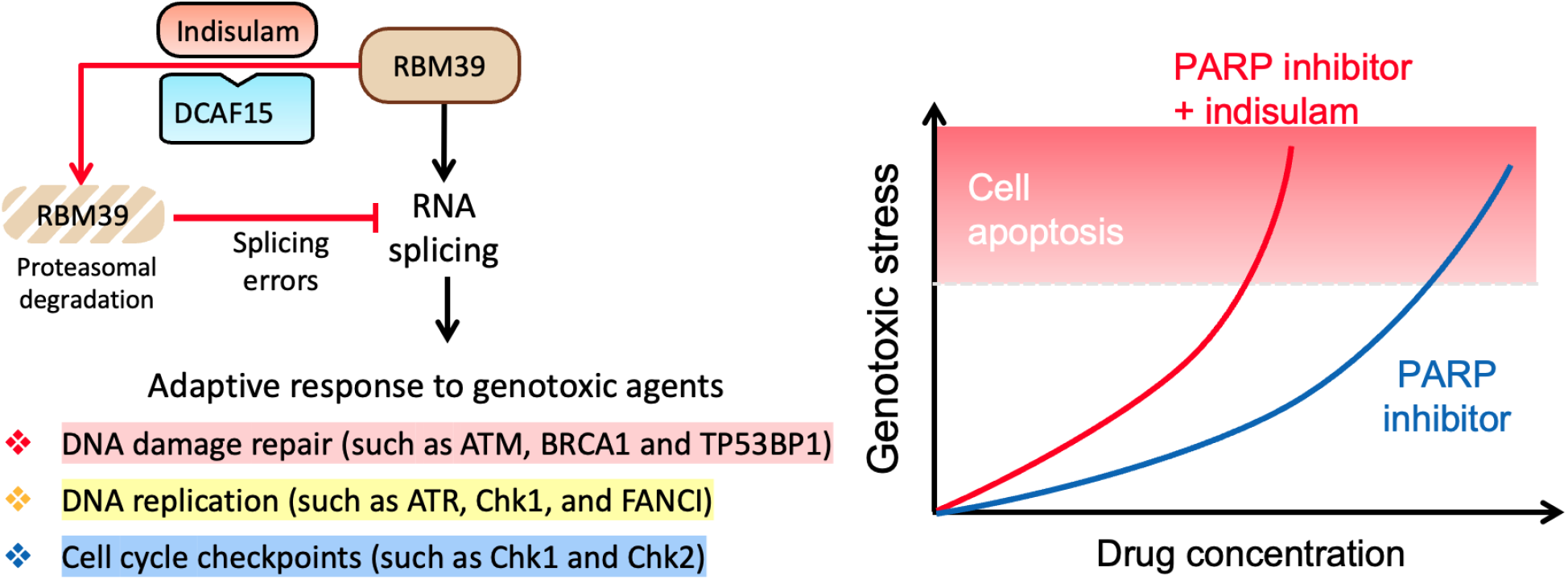

**One Sentence Summary:** We identified a novel drug combination that may improve PARP inhibitor response and benefit a large group of ovarian cancer patients.

## INTRODUCTION

Ovarian cancer is the deadliest gynecological cancer, with a five-year survival rate lower than 50% (*1*). High-grade serous carcinoma (HGSC) is the most lethal subtype of ovarian cancer, accounting for approximately 70% ovarian cancer cases (*2*). The standard therapy for newly diagnosed ovarian cancer involves debulking surgery followed by carboplatin and paclitaxel chemotherapy. However, the disease often relapses with increased chemoresistance (*3*).

Poly-ADP ribose polymerase (PARP) inhibitors have demonstrated significant clinical benefits in ovarian cancer patients with homologous recombination deficiency (HRD) (*4*). HRD can arise from deleterious mutations in *BRCA1/2* or loss of function in other key homologous recombination (HR) components, rendering cells unable to repair double-stranded DNA breaks (DSBs) accurately through HR repair (*5, 6*). By inhibiting PARP1/2-mediated repair of single-stranded DNA breaks (SSBs), PARP inhibitors achieve synthetic lethality in tumors with HRD (*7, 8*). Three PARP inhibitors, olaparib, rucaparib, and niraparib, have been approved for clinical use in subsets of ovarian cancer patients, demonstrating substantial benefits in delaying disease recurrence and improving overall survival (*9-13*). However, approximately half of ovarian cancer patients are HR-proficient and less responsive to PARP inhibitors (*5*). Long-term administration of the drugs also leads to acquired PARP inhibitor resistance and reduced therapeutic response [reviewed in (*14*)]. Additional interventions are therefore needed to improve PARP inhibitor response.

Splicing of precursor messenger RNAs (pre-mRNAs) is a critical regulatory step in gene expression, during which non-coding introns are spliced out from the final transcript. This process is mediated by the spliceosome, a dynamic protein-RNA complex consisting of splicing factors and small nuclear RNAs (*15-17*). Pre-mRNAs of multi-exon genes can be spliced in several ways when alternative splice sites are recognized by splicing factors, generating mRNAs with different exon combinations and subsequently translating to protein isoforms. Alternative RNA splicing diversifies the transcriptome and proteome and is fundamental to cell survival. However, dysregulated alternative splicing can promote tumorigenesis and chemoresistance (*18-22*). In ovarian cancer, splicing factors are frequently upregulated (*23*), and several subsets of alternative splicing events have been associated with disease progression and patient prognosis (*24-30*). Many DNA repair genes are regulated by alternative splicing and targeting spliceosome components have been shown to sensitize cancer cells to PARP inhibitors by disrupting the expression of genes such as *ATM, RAD51* and *CHEK1* (*31-34*).

The RNA-binding protein RBM39 is a transcriptional co-regulator and splicing co-factor, and a promising therapeutic target in cancer [reviewed in (*35*)]. Targeting RBM39 had been challenging until the recent discovery that RBM39 is selectively degraded by a panel of anticancer aryl sulfonamides, indisulam (E7070), E7820, chloroquinoxaline sulphonamide (CQS), and tasisulam (*36, 37*). These drugs act as molecular glues inducing a neo-interaction between RBM39 and DCAF15-associated E3-ubiquitin ligase complex, resulting in polyubiquitination and proteasomal degradation of RBM39 (Fig. 1A) (*36, 37*). Loss of RBM39 leads to transcriptome-wide splicing abnormalities and causes tumor regression in multiple cancer types, including lung cancer (*38*), colorectal cancer (*36, 38*), acute myeloid leukemia (*39*), and neuroblastoma (*34, 40*), but its impact on ovarian cancer has not been studied, to date.

**Fig. 1.**
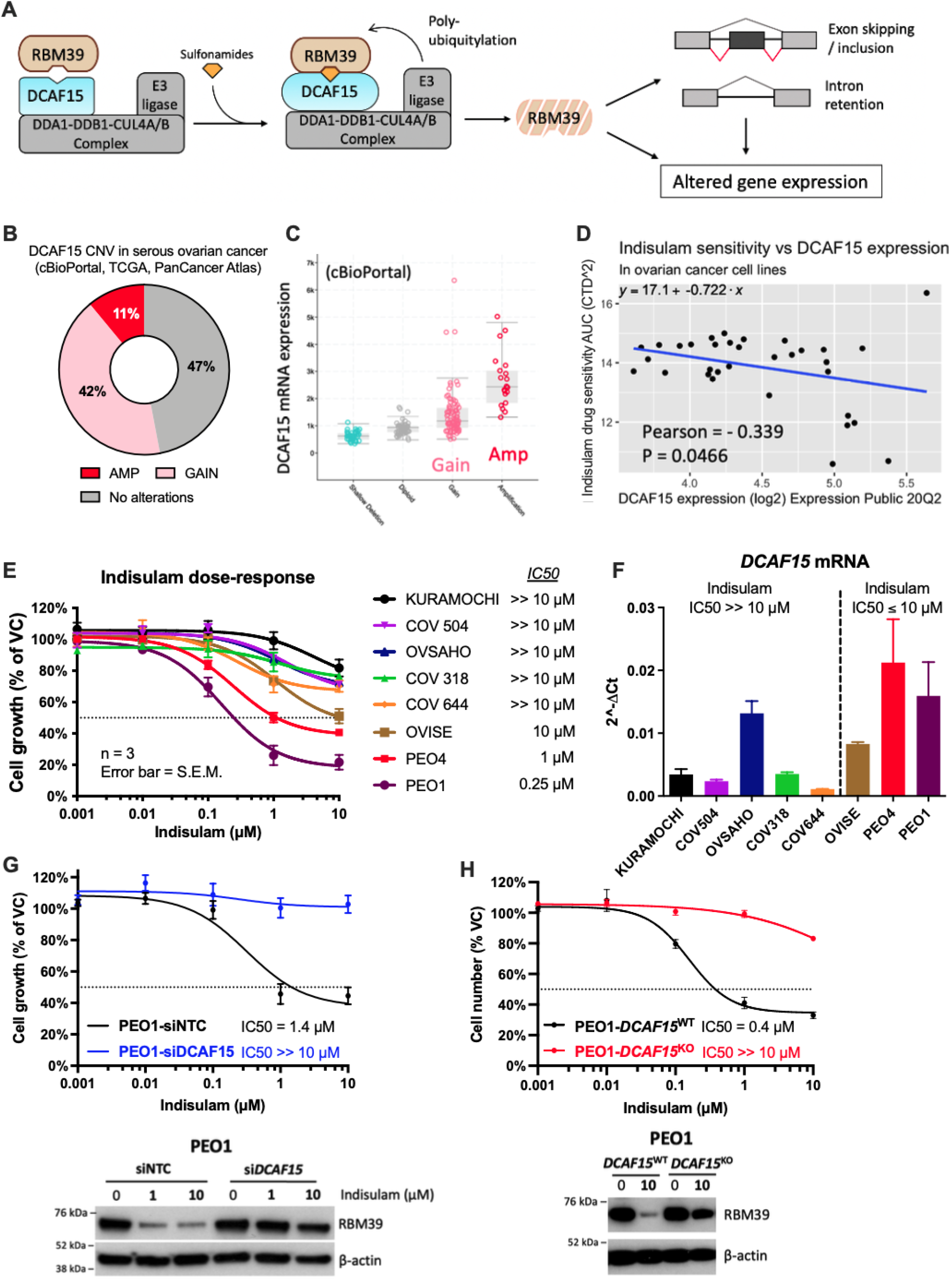
DCAF15 expression correlates to indisulam sensitivity in ovarian cancer cell lines. **(A)** Indisulam degrades RBM39 through a neo-interaction between RBM39 and DCAF15-associated E3 ubiquitin ligase. Loss of RBM39 disrupts the spliceosome and causes widespread splicing abnormalities and altered gene expression. We previously published this diagram (*35*) and it is reused here with permission. **(B)** Copy number variation (CNV) of *DCAF15* in serous ovarian cancer showed frequent amplification (AMP) and gain of copy number (GAIN). Data were extracted from the TCGA PanCancer Atlas on cBioPortal (*41, 42*). **(C)** Amplification or gain of *DCAF15* correlates with increased mRNA expression in serous ovarian cancer. **(D)** Higher *DCAF15* expression was associated with increased sensitivity to indisulam (Area-under-curve, AUC). Data from DepMap portal (https://depmap.org/portal/). **(E)** Human ovarian cancer cell lines were treated with increasing doses of indisulam for 72 hours and cell growth was measured by SRB assay (*n*=3, independent experiments). The concentration causing 50% growth inhibition was defined as IC50. **(F)** The mRNA level of *DCAF15* in ovarian cancer cell lines was quantified using PCR (*n*=3). **(G)** PEO1 cells transfected with DCAF15 siRNA (siDCAF15) or non-targeting control (siNTC) before measuring indisulam sensitivity using the SRB assay (top, *n*=3) and RBM39 degradation (bottom, six hour treatment, *n*=3). **(H)** PEO1 cells were edited with CRISPR-Cas9 to generate *DCAF15* knockout clones (PEO1-*DCAF15*^KO^) and mock editing clones (PEO1-*DCAF15*^WT^). Cell growth (*n*=3) and RBM39 degradation (48h, *n*=3) in these clones were investigated using SRB and western blot, respectively. Dashed lines in (e), (g), and (h) indicate 50% growth inhibition. All data are mean ± SEM

In our recent publication (*40*), we performed RNA-sequencing analysis using IMR-32 neuroblastoma cells treated with indisulam. We detected many aberrant splicing events affecting multiple DNA repair genes, suggesting that RBM39 may regulate the splicing of DNA repair genes. Therefore, we hypothesized that RBM39 has similar functions in ovarian cancer and that targeting RBM39 may disrupt DNA repair pathways and improve PARP inhibitor response. In this study, we explored the therapeutic potential of RBM39 as a cancer target in ovarian cancer. We found that loss of RBM39 causes splicing errors in DNA repair genes in a *DCAF15*-dependent manner, leading to increased sensitivity to DNA damaging agent cisplatin and multiple PARP inhibitors. The combination of indisulam and olaparib caused synergistic growth inhibition and induction of cell apoptosis. The synergy resulted from indisulam exacerbating genotoxic stress from olaparib treatment, as shown by significantly higher γH2AX levels and DNA replication abnormalities. This was associated with suppressed activation of ATM, Chk1, and Chk2, suggesting compromised DNA damage response and replication stress response. Notably, the synergy was observed in multiple ovarian cancer cell lines irrespective of *BRCA1/2* status and PARP inhibitor resistance, suggesting that it may benefit a broad range of patients. Using the ID8 murine HGSC model, we confirmed that indisulam significantly improves the survival benefits of olaparib in animals bearing HR-proficient tumors. Overall, this study provides evidence that targeting RBM39 is a promising strategy for ovarian cancer and could improve the clinical performance of PARP inhibitors.

## RESULTS

### Indisulam causes RBM39 degradation and mis-splicing in ovarian cancer

*In silico* analysis of the Cancer Gene Genome Atlas (TCGA) PanCancer Atlas dataset via cBioPortal (*41, 42*) showed that up to 53% of ovarian high grade serous carcinomas had copy number gain of *DCAF15*, which correlated with upregulated mRNA expression (Fig. 1, B and C). We next investigated the relationship between *DCAF15* mRNA abundance and indisulam sensitivity using the Cancer Target Discovery and Development (CTD^2^) Network data (*43-45*). We found that *DCAF15* expression levels in ovarian cancer cell lines positively correlated with indisulam sensitivity (p = 0.0466), which is consistent with findings in other cancer types (*34, 36, 37, 39, 40*) (Fig. 1D). This association was observed *in vitro* in ovarian cancer cells: cells with higher *DCAF15* expression were more sensitive to indisulam (Fig. 1, E and F), and indisulam caused a dose-dependent degradation of RBM39 (Fig. 1E and S1). Knockdown (via siRNA) or knockout (via CRISPR-Cas9) of *DCAF15* in PEO1 cells conferred resistance to indisulam and rescued indisulam-mediated RBM39 degradation (Fig. 1G-H and S1A), implying that DCAF15-mediated RBM39 degradation is necessary for efficacy. Interestingly, even in the most resistant cell line, KURAMOCHI, DCAF15 levels were sufficient to degrade RBM39 in response to indisulam treatment (Fig. S1B). This indicates that cell dependency on RBM39 and not just DCAF15 levels determine response to indisulam. Taken together, our data demonstrate that indisulam inhibits ovarian cancer proliferation by depleting RBM39, and that DCAF15 expression is necessary but not sufficient to predict indisulam cytotoxicity.

### RBM39 regulates the splicing of DNA damage repair genes

Transcriptome-wide characterization of alternative splicing events after RBM39 depletion has been investigated in multiple cancer types but not in ovarian cancer. Although there could be cancer-specific differences, we speculate that RBM39 modulates a robust RNA splicing signature across various cancer types. To identify common alternative splicing targets of RBM39, we analyzed 11 publicly available RNA sequencing datasets with alternative splicing analysis in RBM39-depleted cancer cells. These include five datasets in neuroblastoma cells (BE2C and SKNAS) (*34*), three melanoma cells (501-MEL-ES, A375, and SK-MEL-239) (*46*), two in acute myeloid leukemia cells (MOLM-13 and THP-1) (*39*), and one in MCF-7 breast cancer cells (*47*) (Fig. S2, A and B). These datasets were analyzed in combination with our recently published splicing analysis in neuroblastoma (*40*). We found that, upon RBM39 depletion, exon skipping events in 1774 genes and intron retention in 629 genes were repeatedly detected across cancer types (Table S1). Pathway enrichment analysis revealed that cell cycle and DNA repair pathways are preferential splicing targets of RBM39, indicating a consistent role of RBM39 in modulating cell cycle progression and DNA damage repair response (Fig. S2C).

To investigate the impact of RBM39 depletion with indisulam in ovarian cancer, we performed RNA sequencing and alternative splicing analysis using SpliceFisher and rMATS algorithms on KURAMOCHI cells. The analysis revealed that indisulam predominantly induced exon skipping or intron retention events (Fig. 2A, Fig. S2D). These genes were highly enriched for pathways related to RNA metabolism, cell cycle, and DNA damage repair (Fig. 2B, Fig. S2D). Notably, indisulam caused aberrant splicing events in many DNA repair genes, such as *ATM, ATR*, Fanconi Anemia genes (*FANCI*), HR genes (*BRCA1* and *RAD51D*), and non-homologous recombination (i.e. non-homologous end joining, NHEJ) (*TP53BP1* and *RIF1*) (Fig. 2, C and D). Alternative splicing of these DNA repair genes is also affected in other cancer types (Fig. 2C), suggesting a conserved role for RBM39 in regulating the alternative splicing of key players in the response to DNA damage.

**Fig. 2.**
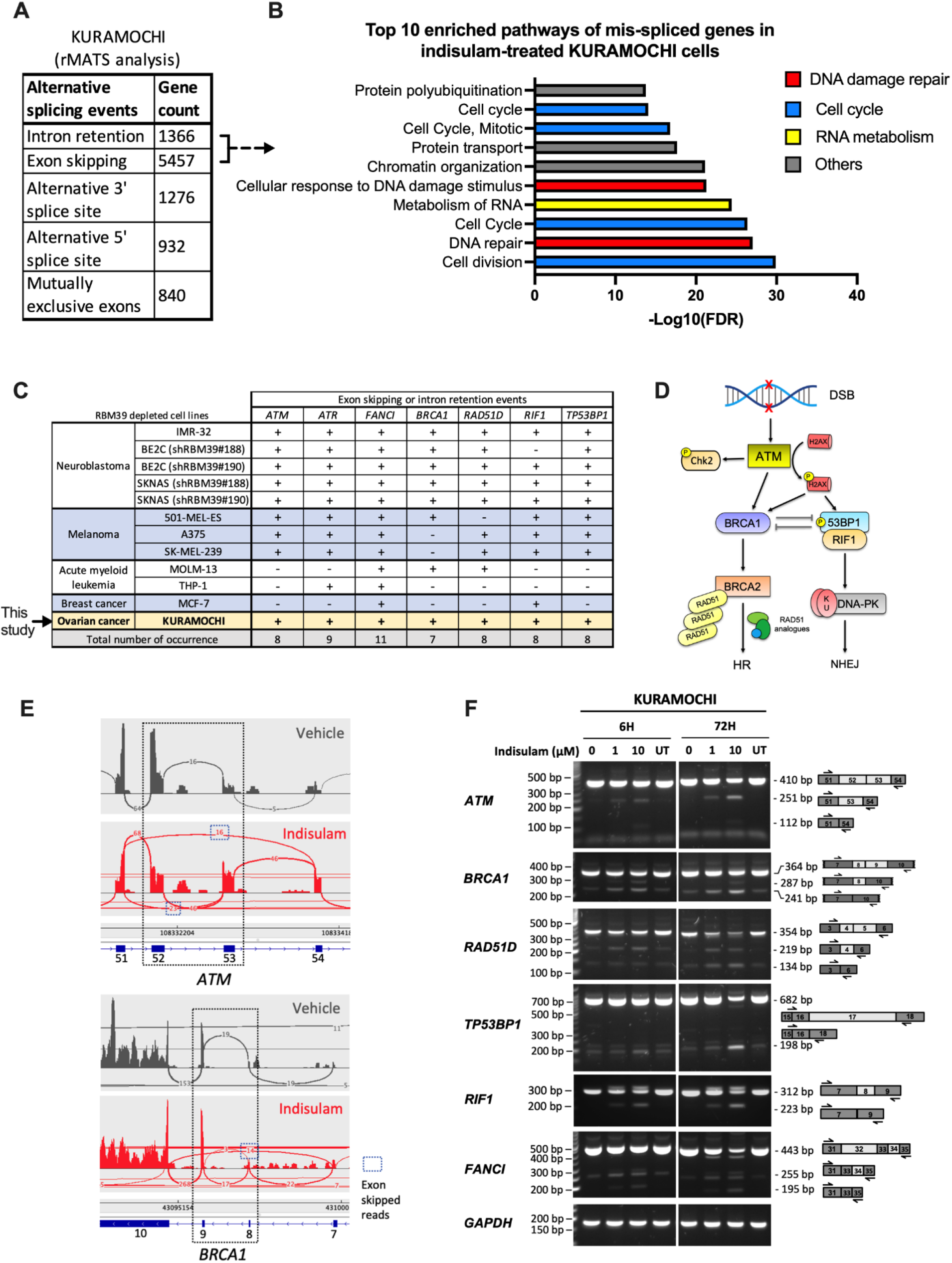
RBM39 depletion causes splicing errors in DNA damage repair genes. **(A)** Alternative splicing events as determined by rMATS in KURAMOCHI cells following exposure to indisulam (24h, *n*=3 independent experiments). **(B)** Pathway enrichment analysis (FDR<0.1) on intron-retained or exon-skipped genes **(C)** Meta-analysis of 11 alternative splicing datasets in RBM39-depleted cells, including our datasets in KURAMOCHI and IMR-32 (*40*). Common key damage repair genes are depicted, “+”, genes exon skipped or intron retained; “-”, no exon skipping or intron retention events detected. **(D)** A simplified diagram highlighting the role of ATM, BRCA1, RAD51 analogous, 53BP1, and RIF1 in DSB repair. **(E)** Sashimi plots of *ATM* (skipping of exon 52/53) and *BRCA1* (skipping of exon 8/9) in KURAMOCHI following indisulam exposure. **(F)** PCR validation of indisulam-induced exon skipping events in DNA repair genes (representative image of *n*=3). *GAPDH* was included as a control. UT, untreated.

To validate the changes in splicing, we used PCR with custom-made primers flanking the affected exons of *ATM, BRCA1, RAD51D, TP53BP1, RIF1*, and *FANCI*, which permits the detection of exon skipping events. PCR analysis showed that indisulam exposure results in aberrant splicing in these genes (Fig. 2E, 2F, and Fig. S3), potentially leading to truncated variants that are unlikely to translate to functional proteins due to frameshifts or deletions in functional domains. For example, skipping of exon 52 and 53 in *ATM* mRNAs are likely to disrupt the catalytic activity of ATM due to deletions (aa. 2544-2642) in the kinase domain (*48, 49*). The splicing variants of *ATM* lacking exon 53 (139 bp), *BRCA1* lacking exon 9 (77 bp), and *TP53BP1* lacking exon 17 (484 bp) are expected to produce frameshifts and premature stop codons, leading to nonsense-mediated decay (*50*).

Overall, our findings demonstrate that RBM39 depletion modulates the splicing of DNA repair genes across multiple cancer types, including ovarian cancer. Targeting RBM39 may create vulnerabilities in cancer cells to anticancer drugs that increase genotoxic stress.

### Indisulam synergizes with cisplatin and PARP inhibitors in ovarian cancer

Since indisulam induces splicing errors in DNA damage repair genes, we speculated that this may disrupt DNA damage response and render ovarian cancer cells susceptible to DNA damaging agents such as cisplatin and PARP inhibitors. KURAMOCHI and OVSAHO cells were used to test this hypothesis because they are representative HGSC cell lines (*51*), and are intrinsically resistant to cisplatin and PARP inhibitors compared to other HGSC models with loss of BRCA1/2 (i.e. PEO1) (Fig. 3A, 3B, and S4). Combination index (CI) values were calculated according to the Chou-Talalay method (*52*). Combination of indisulam and cisplatin synergistically inhibited growth in KURAMOCHI and OVSAHO cells compared to the monotherapies (Fig. 3C upper panels). Furthermore, indisulam significantly elevated caspase-3/7-dependent apoptosis when combined with cisplatin (Fig. 3C lower panels).

**Fig. 3.**
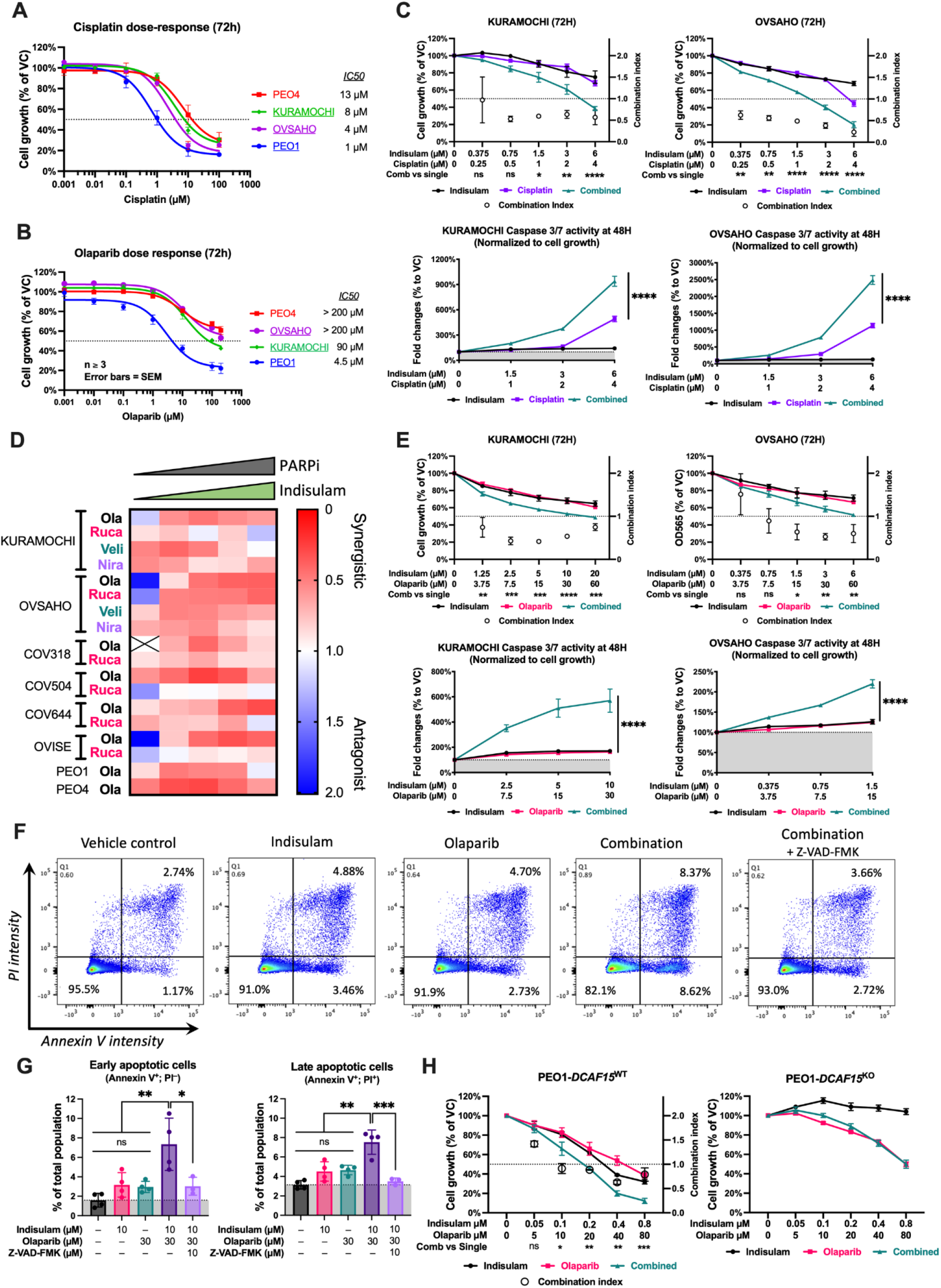
Indisulam improves cisplatin and PARP inhibitor response in ovarian cancer cells. **(A-B)** Growth assay (SRB) of four ovarian HGSC cell lines (PEO1, PEO4, KURAMOCHI, and OVSAHO treated with increasing doses of cisplatin (a) and olaparib (b) for 72 hours (*n*=3 independent experiments, PEO4; cisplatin, *n*=2). Cell lines with *BRCA2* mutation or deletions are underlined. **(C)** Cell growth and apoptosis in KURAMOCHI and OVSAHO cells treated with indicated doses of indisulam and cisplatin either alone or in combination (*n*=3 independent experiments). Open circles indicate Combination Index (CI) values (right y-axis). **(D)** Heatmap of CI values of indisulam in combination with indicated PARP inhibitors in a panel of human ovarian cancer cell lines. **(E)** Cell growth and apoptosis of KURAMOCHI and OVSAHO cells after treatment of indisulam, olaparib, or combination (*n*=3 independent experiments). Open circles indicate CI values (right y-axis) **(F)** KURAMOCHI cells exposed to indisulam and olaparib individually or in combination (with or without 10 μM Z-VAD-FMK) for 48 hours. Apoptosis was analyzed by flow cytometry using dual labeling of Annexin V and propidium iodide (PI) (n>3 independent experiments). **(G)** Quantification of early apoptotic and late apoptotic cells from flow cytometry analysis in (f). Ordinary one-way ANOVA analysis and Tukey’s multiple comparison tests were used for statistical analysis. **(H)** Growth assay of PEO1-*DCAF15*^WT^ and PEO1-*DCAF15*^KO^ cells treated with indisulam and olaparib either alone or in combination for 72 hours (*n*=3 independent experiments) Open circles indicate CI values (right y-axis). Statistical significance between drug combination and individual treatment (c, e, and h) was determined by two-way ANOVA and Dunnett’s multiple comparisons test. ns=not significant, *p<0.05, **p<0.01, ***p<0.001, ****p<0.0001. CI values lower than 1 indicate synergism and CI values larger than 1 indicate antagonism (error bars indicate ±SD). Data in indicate mean ± SEM (a, b, c, e, h) or ± SD (f).

Toxicity is a major concern for combination therapies with platinum agents and combining indisulam with PARP inhibitors could be a safer strategy. To test whether indisulam improves PARP inhibitor response in ovarian cancer cells, we combined indisulam with four clinical PARP inhibitors olaparib, rucaparib, veliparib, or niraparib in eight human ovarian cancer cell lines (Fig. 3D and S5). Synergistic interactions were observed in almost all drug combinations and all cell lines regardless of drug sensitivity, *BRCA* mutational status, or PARP trapping capacity (Fig. 3D, S4, and S5). Subsequent cell growth and apoptosis assays validated that combination therapy of indisulam and olaparib caused synergistic growth inhibition and cell apoptosis in KURAMOCHI and OVSAHO cells (Fig. 3E), even though these two cell lines are relatively resistant to both indisulam and olaparib monotherapies. Flow cytometry analysis further demonstrated that the Annexin V+/PI+ population induced by the drug combination could be rescued by the pan-caspase inhibitor Z-VAD-FMK, confirming apoptosis as the major form of cell death (Fig. 3, F and G). Meanwhile, a short-term exposure (48 or 72 hours) to indisulam and olaparib in combination was sufficient to induce a synergistic and long-lasting inhibitory effect on HGSC colony growth (at least nine days after removing drug, Fig. S6), a desirable characteristic for future clinical translation.

These observations are in line with our hypothesis that indisulam compromises DNA repair pathways through interference of RNA splicing and is able to potentiate the impact of clinical DNA damaging agents.

Exon skipping events of DNA repair genes, such as *ATM* and *TP53BP1*, were confirmed in cells treated with indisulam alone and in combination with olaparib (Fig. S7). CRISPR-Cas9 knockout of DCAF15 completely abolished the synergistic interaction between indisulam and olaparib, suggesting that DCAF15-mediated RBM39 degradation and subsequent splicing changes are necessary for the synergy (Fig. 3H). This is consistent with that the fact that E7820, another molecular glue degrading RBM39 via DCAF15, also acts in synergy with olaparib (Fig. S8).

Interestingly, the synergistic interaction between indisulam and olaparib seems independent of HRD status as it was observed in both HR-deficient PEO1 cells and a patient-matched HR-proficient PEO4 cells (Fig. S9A). The same trend was also seen in an isogenic pair of murine HGSC cell lines: ID8-*Trp53*^*–/–*^ and ID8-*Trp53*^*–/–*^*;Brca2*^*–/–*^ (Fig. S9B)(*53*).

Taken together, our findings demonstrate that combining indisulam or other RBM39-targeting molecular glues with PARP inhibitor treatment is a promising strategy to ovarian treat HGSC, which could benefit a wide range of patients regardless of their *BRCA1/2* status, or PARP inhibitor regime. Importantly, ovarian HGSC with intrinsic and acquired resistance could potentially be re-sensitized to PARPi when combined with RBM39 degraders.

### Indisulam treatment exacerbates genotoxic stress caused by Olaparib

PARP inhibitors increase genotoxic stress by inducing DNA damage (SSBs and DSBs) and promote replication fork abnormalities (*54*). The ATM-Chk2-mediated DNA damage repair and the ATR-Chk1-mediated replication stress response are two mechanisms that promote cancer cell survival in response to PARP inhibitor treatment (*55-58*).

Since indisulam-mediated RBM39 degradation induces aberrant splicing errors in ATM and downstream DNA repair genes, we hypothesized that indisulam sensitizes ovarian cancer cells to PARP inhibitors by suppressing ATM-mediated DNA damage response. Western blot analysis in KURAMOCHI cells revealed that indisulam decreased ATM protein expression and suppressed olaparib-induced phosphorylation of ATM (pATM-S1981) and Chk2 (pChk2-T68) (*59*), indicating that indisulam impaired the ability to mount a DNA damage response (Fig. 4A). We assessed the level of DNA damage with phospho-H2AX (γH2AX) which increases in response to DSBs (*60, 61*) and stalled replication forks (*61-66*). Western blot data showed that, whilst indisulam and olaparib monotherapies had a minor impact on γH2AX expression, combination therapy led to a substantial increase in γH2AX levels, suggesting a synergistic induction of DSBs or replication stress (Fig. 4A). Similarly, immunofluorescence demonstrated that indisulam and olaparib potentiate γH2AX foci formation (Fig. 4B and S10A), which was also observed when indisulam was combined with other PARP inhibitors (Fig. S10B-G).

**Fig. 4.**
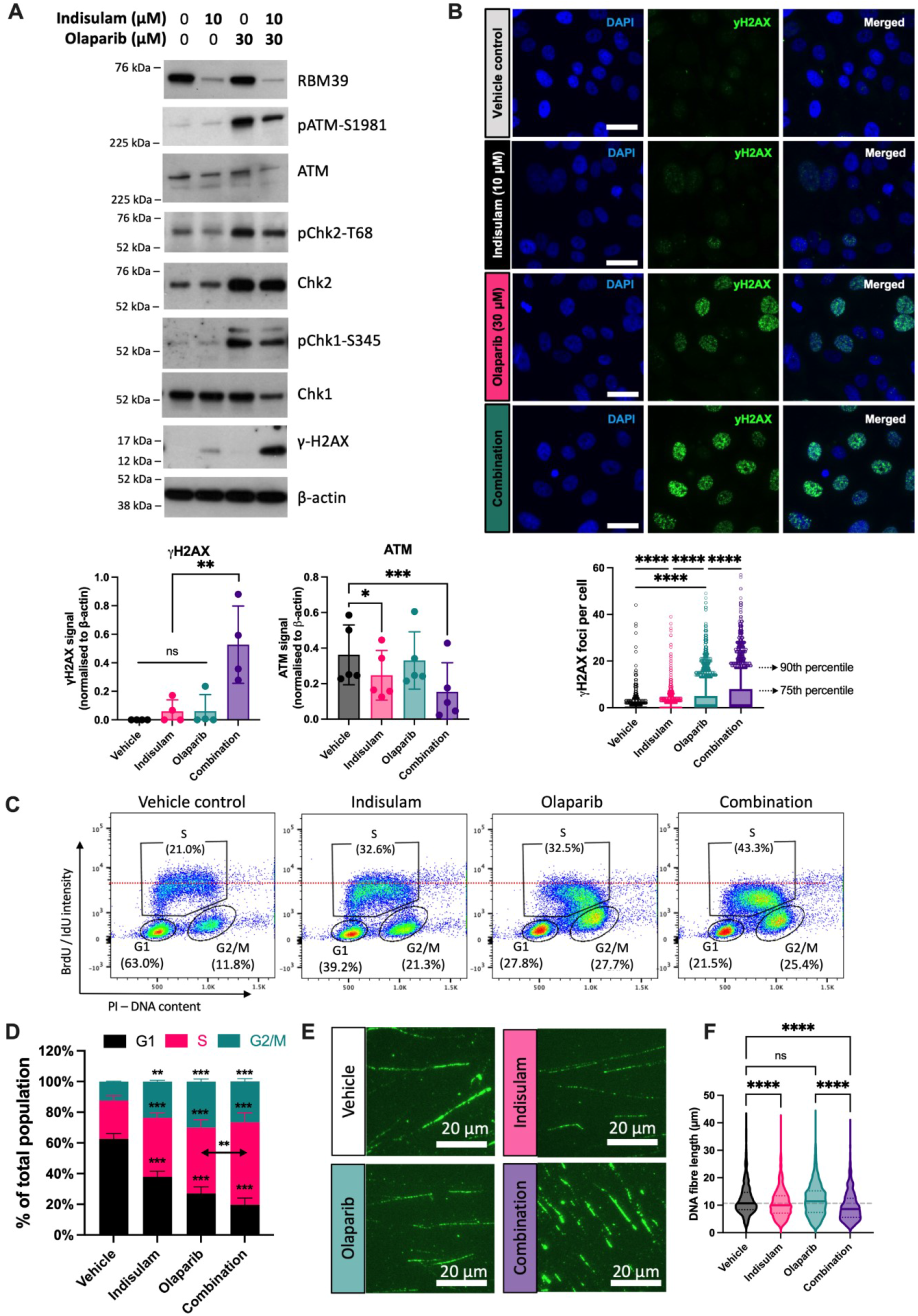
Indisulam inhibits ATM expression and exacerbates genotoxic stress caused by PARP inhibitors. **(A)** Western blot analysis (top) and densitometry analysis (bottom) of KURAMOCHI cells after 48 hour treatment of indicated drugs (representative blot from n>3 independent experiments). One-way ANOVA and Dunnett’s multiple comparison test were used to determine the statistical significance. **(B)** Immunofluorescence of KURAMOCHI cells treated with indisulam or olaparib as monotherapies or in combination for 24 hours (top panel, representative image). Analysis of γH2AX foci from three independent experiments. (bottom panel, whiskers indicate 10^th^ and 90^th^ percentiles). Size bar, 50 μm. **(C)** Flow cytometry-based cell cycle profiling of KURAMOCHI cells after 48 hour treatment of 10 μM indisulam, 30 μM olaparib, or the combination. Cellular DNA content was labelled with propidium iodide (PI) and replicating cells were labelled with nucleotide analogs 5-Bromo-2’Deoxyuridine (BrdU) and 5-Iodo-2-deoxyuridine (IdU). **(D)** Analysis of cell cycle analysis (*n*=3 independent experiments). Two-way ANOVA and Tukey’s multiple comparisons test were used to determine statistical significance among groups. **(E)** DNA fiber analysis quantifying replication fork behaviors in KURAMOCHI cells after 48-hour exposure to indisulam, olaparib, or combination. Representative fields are shown. Scale bar, 20 μm. **(F)** Violin plots (with median and interquartile range) of DNA fiber length measurements (at least 300 fibers from 2 independent experiments). Kruskal-Wallis test and Dunn’s multiple comparisons test were used to determine statistical significance. ns = not significant, *p<0.05, **p<0.01, ***p<0.001, ****p<0.0001. All data are mean ±SD.

Flow cytometry analysis revealed KURAMOCHI cells treated with the combination of indisulam and olaparib were more likely to arrest in the S phase (mean = 53.8%, p < 0.01) than monotherapies of indisulam (mean = 38.5%) and olaparib (mean = 42.9%) (Fig. 4, C and D). Interestingly, cells treated with the combination therapy had the least nucleotide incorporation (mean = 47% of vehicle control; Fig. S12A), implying reduced DNA synthesis. This is in line with a reduction in levels of total and phosphorylated (S345) Chk1 (Fig. 4A). Chk1 safeguards DNA replication by restarting stalled replication forks and inhibits unscheduled new origin firing (*67-71*). Therefore, we hypothesized that the combination of indisulam and olaparib might cause DNA replication abnormalities. To this end, we examined DNA replication fork behaviors using DNA fiber analysis. Whilst indisulam and olaparib alone had minor to moderate impacts on the length of newly synthesized DNA (DNA fibers), their combination led to significantly shorter DNA fibers, indicating increased fork stalling or reduced fork restarting (Fig. 4, E and F). Meanwhile, the combination therapy led to a profound increase in new origin firing (Fig. S11, B and C), which is consistent with the effect of Chk1 inhibition (*67, 72, 73*).

Taken together, our data indicate that indisulam increases PARP inhibitor response by exacerbating genotoxic stress, with concomitant suppression of the ATM-Chk2-mediated DNA damage repair and Chk1-modulated replication stress response.

### Combination of indisulam and olaparib demonstrates significant survival benefits *in vivo*

Next, we evaluated the combination of indisulam and olaparib in a well-characterized mouse model of ovarian HGSC (ID8) (*53, 74*). This model recapitulates the process of peritoneal metastasis and development of ascites, observed in ovarian HGSC patients, and mimics the tumor immune microenvironment which is advantageous over traditional xenografts (Fig. 5A) (*53*). Given that indisulam synergized with olaparib in ID8 cells irrespective of *Brca2* status *in vitro* (Fig. S9), we hypothesized that the drug combination would be effective for HR-proficient tumors. The ID8-*Trp53*^*–/–*^ clone was therefore selected for *in vivo* analysis.

**Fig. 5.**
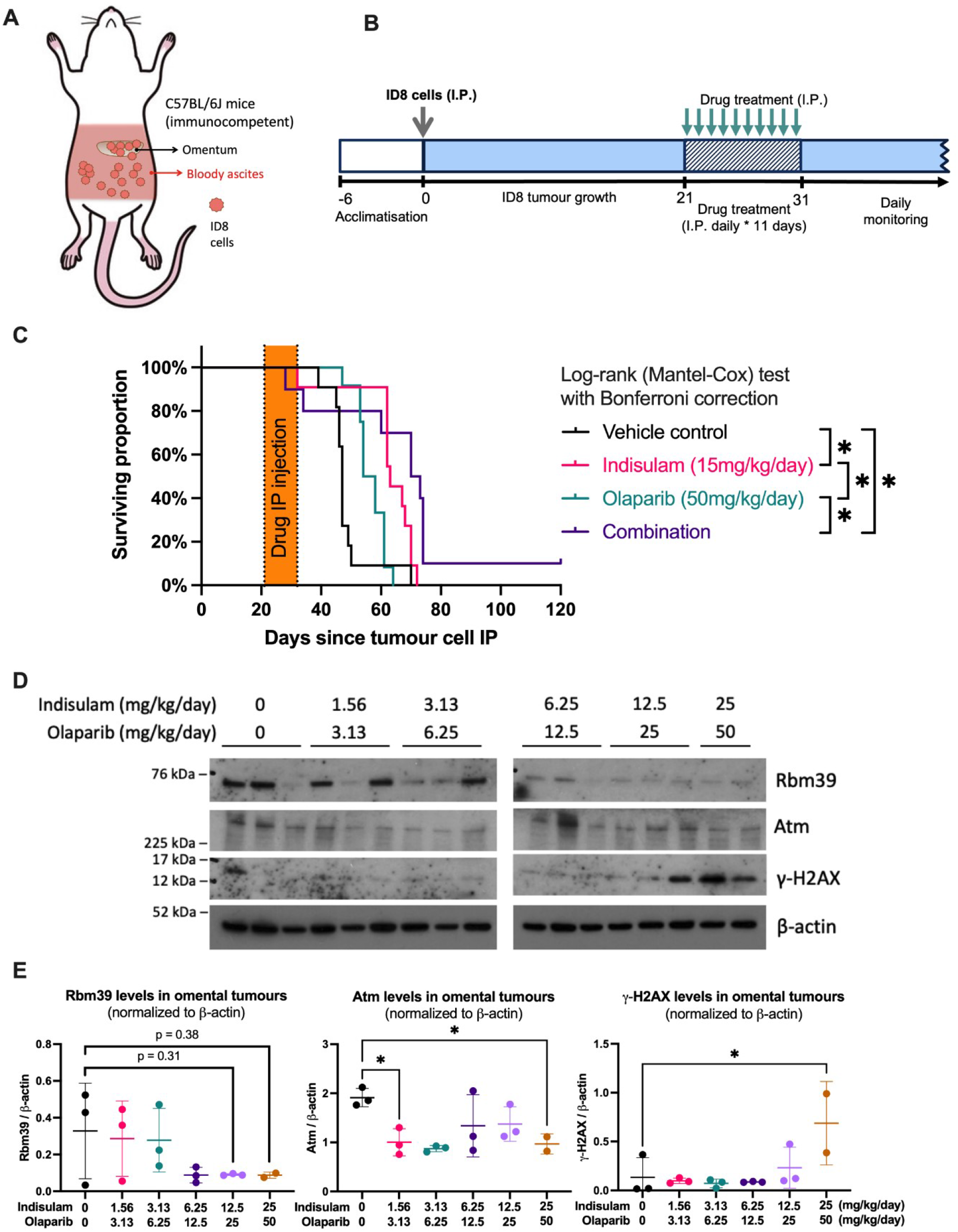
Indisulam improves olaparib response *in vivo*. **(A-B)** Schematic of ID8 murine HGSC model with the formation of peritoneal metastasis and ascites and survival analysis design. **(C)** Kaplan-Meier survival curves of mice bearing ID8 tumors after indicated treatments (*n*=11 for vehicle control, *n*=11 for indisulam, *n*=12 for olaparib, and *n*=10 for drug combination). Statistical significance was calculated using the log-rank (Mantel-Cox) test and Bonferroni multiple comparisons test. *p<0.0083 (Bonferroni-corrected threshold). **(D)** Western blot assessment of omental tumor samples collected from mice with indicated drug doses daily for seven days (or five days in the case of the highest dose). **(E)** Densitometry analysis of Rbm39, Atm, and γH2AX bands intensities in (d). Error bar = SD. One-way ANOVA and Dunnett’s multiple comparisons test were used for statistical analysis. *p<0.05.

Pilot studies were conducted to determine maximal tolerable doses and optimal dosing schedules of indisulam and olaparib; additional adverse effects were not observed with combination therapy (Table S2). Daily intraperitoneal administration of up to 15 mg/kg of indisulam and 50 mg/kg of olaparib for 11 days were well tolerated and used for survival analyses (Fig. 5B). In comparison with mice receiving vehicle control (median survival = 47 days), olaparib treatment alone only offered moderate survival benefits (median survival = 56 days, p=0.0206) whilst indisulam alone improved the median survival to 65 days (p=0.0063) (Fig. 5C). However, the combination of indisulam and olaparib resulted in longest survival and was significantly more effective than olaparib monotherapy (median survival = 73 days, p=0.0041). Interestingly, one of the mice treated with the drug combination survived till end study (120 days) and showed complete tumor regression (Fig. 5C).

Western blot analysis of the omental tumor deposits exposed to combination therapy indicated a dose-dependent degradation of Rbm39 confirming the on-target effect of indisulam (Fig. 5, D and E). Downregulation of Atm and accumulation of γH2AX were also observed in tumor tissues following simultaneous administration of indisulam and olaparib, which is consistent with *in vitro* observations (Fig. 5, D and E). In summary, these data demonstrate that indisulam improves olaparib response in HR-proficient ovarian cancer *in vivo*, highlighting the promising therapeutic value of indisulam to expand the clinical use of PARP inhibitors in ovarian HGSC.

## DISCUSSION

Ovarian HGSC is a leading cause of gynecologic cancer-related death, with limited options for targeted therapies (*2, 75-77*). Routine administration of PARP inhibitors has led to the emergence of drug resistance, despite promising benefits of delaying disease relapse and improving the overall survival of ovarian cancer patients (*78*). PARP inhibitor resistance can occur in multiple ways. Reversion mutations that restore HR proficiency are the most common mechanism in the clinic (*14*). Other resistance mechanisms include decreased PARP trapping, upregulated drug efflux, and stabilization of stalled DNA replication forks (*14*). Our systematic literature and transcriptomic analysis uncovered the role of RBM39 in coordinating RNA splicing of cell cycle and DNA damage repair genes in various cancer cells. Using a variety of functional assays, we show that RBM39-degrading molecular glues such as indisulam can be repurposed as a therapeutic agent to improve PARP inhibitor response *in vitro* and *in vivo*. Importantly, a synergistic interaction between indisulam and various PARP inhibitors was observed in multiple ovarian HGSC cell lines *in vitro*, including BRCA-mutated and PARP inhibitor-sensitive cell lines (PEO1 and ID8-*Trp53*^*– /–*^*;Brca2*^*–/–*^) and PARP inhibitor-resistant cell lines (KURAMOCHI, OVSAHO, PEO4, COV318, COV504, COV644, and ID8*-Trp53*^*–/–*^). Synergy occurs in a DCAF15-dependent manner, excluding the involvement of previously reported off-target effects of indisulam (*40, 79-81*). Notably, in the *Brca1/2*-wildtype ID8 model (ID8*-Trp53*^*–/–*^), the combination of indisulam and olaparib offered greater survival benefits than either monotherapy, demonstrating that indisulam improves PARP inhibitor response in mice with PARP inhibitor-resistant ovarian tumours. Further, our findings are consistent with a recent CRISPR-dropout screen, where sgRBM39 was significantly depleted upon olaparib treatment or PARP1-knockout (*82*).

Mechanistically, indisulam disrupts RBM39-mediated splicing of cell cycle, DNA damage repair, and DNA replication genes, leading to reduced protein expression and a defective adaptive response to genotoxic stress caused by PARP inhibitors (Fig. 6). Indisulam downregulated ATM expression at protein levels, likely due to splicing errors causing frameshifts. Indisulam-mediated ATM downregulation has also been documented in neuroblastoma cells (*34*), but the authors did not investigate the functional consequences. Indisulam suppresses olaparib-induced activation of ATM-Chk2-mediated DNA damage response and Chk1-mediated replication stress response, associated γH2AX accumulation, increased replication fork abnormalities, cell cycle arrest in the S phase, and cell apoptosis. These phenotypes are similar to those observed with combination of PARPi and ATM inhibitors (*83, 84*) or inhibitors targeting Chk1 or Chk2 (*85-87*). As ATM is upstream of BRCA1/2-mediated HR repair, our model may explain why indisulam synergizes with PARP inhibitors regardless of *BRCA1*/2 status.

**Fig. 6.**
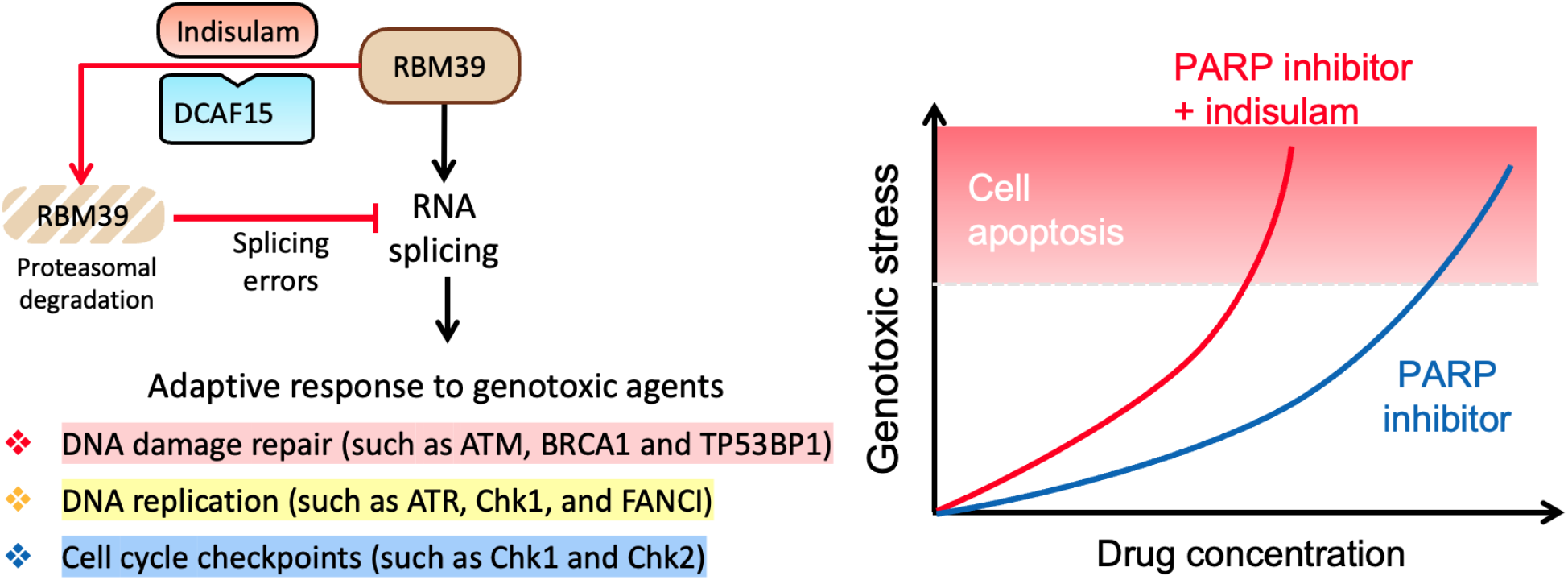
Targeting RBM39 exacerbates genotoxic stress caused by PARP inhibitors. Data from this study suggest a model that RBM39 mediates appropriate splicing of many genes in DNA damage repair, DNA replication, and cell cycle progression, thereby interfering with the adaptive response to genotoxic agents. Depletion of RBM39 with indisulam renders cells susceptible to genotoxic stress, leading to accelerated genotoxic stress accumulation, cell apoptosis, and increased sensitivity to PARP inhibitors.

One interesting finding is that only the combination of indisulam and olaparib, but not monotherapies, downregulate Chk1 expression, suggesting an intriguing interaction between these two drugs. PARP1 has been implicated in pre-mRNA splicing (*88-91*), and it is possible that simultaneous disruption of PARP1- and RBM39-mediated RNA splicing affected the expression of genes such as Chk1. Meanwhile, RBM39 may interact directly or indirectly (through other splicing factors) with PARP1 as PARP1 was co-immunoprecipitated with RBM39 (*39*), and both interact with the core splicing factor SF3B1 (*88, 92*).

Indisulam was discovered as a cell cycle inhibitor in the late 1990s (*93*), but its mechanism of action was only revealed by two seminal studies in 2017 (*36, 37*). Aryl sulfonamides like indisulam are now recognized as a novel class of molecular glues that induces rapid depletion of RBM39 through DCAF15-mediated proteasomal degradation, resulting in splicing defects in over 3000 genes (*36, 37, 39, 40, 94*) and tumor regression in multiple animal models (*36, 38-40*). Indisulam and three other anticancer aryl sulfonamides have been tested in at least 47 clinical trials, but only one clinical trial involved patients with ovarian cancer [reviewed in (*35*)]. Monotherapy of aryl sulfonamides demonstrated minor to moderate clinical benefits, and most of them failed to progress beyond a phase II trial (*35*). Although some tumors may be sensitive to RBM39 depletion alone (like, neuroblastoma (*34, 40*)), here we demonstrate a rational case for the combination therapy of aryl sulfonamides and DNA damaging agents like platinum and PARPi in ovarian HGSC. Although DCAF15 is necessary for RBM39-mediated cytotoxicity and levels of *DCAF15* correlate to sensitivity *in vitro*, expression of *DCAF15* did not predict sensitivity to indisulam in AML patient samples (*98*). This highlights the need for careful consideration for the use of DCAF15 as a clinical biomarker and the potential importance of additional biomarkers in future development. Nonetheless, the availability of drug pharmacodynamic, pharmacokinetic and safety data in patients for both aryl sulfonamides and PARP inhibitors should aid rapid translation of such combinations to clinical studies.

It is important to note that we did observe some adverse effects for indisulam in our PD trial study at a previous reported dose of 25 mg/kg/day (*36, 39, 95, 96*). One possible explanation is that previous studies mostly used severely immunodeficient mice (*36, 38, 96*), while our trial mice with intact immune systems. Indisulam treatment has been associated with the induction of immune response due to splicing variants-derived neoantigens and altered DCAF15 activities (*46, 97*). Hence, we lowered indisulam to 15mg/kg/day for our combination survival trial and found no evidence that indisulam causes extra toxicities when combined with olaparib at this dose.

The splicing errors that occur after RBM39 depletion are robust across various lineages and targets enriched for cell cycle, cell replication and DNA damage repair. Our study focused on two key signaling pathways that respond to DNA damage (ATM) and cell cycle checkpoints (Chk1/2), however future studies warrant investigation of other mis-spliced genes such as *BRCA1, RAD51D, TP53BP1, RIF1* or *FANCI*. Additionally, RBM39 is a transcription co-activator besides being a splicing factor (*35*). It may be conceivable that the some of the effects of aryl sulfonamides also involves transcriptional changes in genes that do not experience splicing errors but are under transcriptional co-regulation by RBM39.

In summary, this study provides strong evidence that combining RBM39 protein degraders and PARP inhibitors is an effective therapeutic strategy for ovarian HGSC. As dosing schedules of indisulam and olaparib have been optimized in the clinic, a clinical trial could be readily designed and conducted to evaluate the efficacy of such combination therapy in patients. Future studies on RBM39 dependency and its involvement in DNA damage repair pathways and replication stress may reveal novel biomarkers besides DCAF15 that can predict patient response to RBM39-targeted therapies. We expect the benefit of this combination therapy can extend to other cancers where DNA damaging agents and PARPi are approved clinically, such as breast cancer and prostate cancer.

## MATERIALS AND METHODS

### Study design

The main objectives of this study were to investigate the therapeutic value of RBM39 as a cancer target in ovarian cancer cells and whether targeting RBM39 improves PARP inhibitor response. We quantified the synergistic potential between drug pairs using the commonly used method: Chou-Talalay method. KURAMOCHI and OVSAHO cells were selected for most functional assays and mechanistic studies because they were two cell lines genetically similar to human HGSC and exhibited intrinsic PARP inhibitor resistance despite the loss of *BRCA2* expression. The syngeneic ID8 murine HGSC model was chosen for the *in vivo* survival analysis because it provides an immune-proficient environment and recapitulates the process of peritoneal metastasis in patients. For quantitative *in vitro* assays, at least three biological repeats were performed, and for qualitative *in vitro* assays, at least one biological repeat was conducted with different conditions (distinct time points, concentrations, or drug pairs). The sample size for animal studies was determined following guidelines published by Charan and Kantharia (*98*). Pilot experiments with minimal sample sizes were conducted to identify optimal dosing schedules. The survival analysis was conducted in two rounds, and data from the first round of the study were used to determine the minimal sample size for the entire survival analysis. Humane endpoints were assessed according to Protocol 6 detailed in the animal study Project License (PA780D61A). Mice were randomized for drug treatment, and researchers were blinded from treatment conditions during the survival analysis.

### *In vivo* experiments

Animal studies were designed and conducted in accordance with the UK Animals (Scientific Procedures) Act 1986. Mice were kept at the Central Biomedical Services (CBS), Imperial College London. All individuals involved were Home Office Personal License holders working under the Project License (PA780D61A). C57BL/6 female mice aged between 6-7 weeks were purchased from the Charles River Laboratory (UK). Newly arrived mice were acclimatized for seven days before regulated procedures. Then, five million ID8-*Trp53*^−/–^ cells were injected intraperitoneally (I.P.) into each mouse and allowed to proliferate for at least 21 days before drug treatments.

Drug solutions of vehicle control (3.5% DMSO and 6.5% Tween-80 in saline) indisulam (Sigma-Aldrich, #SML1225), or olaparib (Selleckchem, #S1060) were prepared for *in vivo* administration (*36, 38, 40*). Left and right abdomen were alternated for I.P. injections (11 days) to minimize animal discomfort. For pharmacodynamic experiments, mice were randomly enrolled into 4 treatment groups (n=3 mice per group) and were culled within 48 hours after the last dosing. For survival studies, mice (n=11 for vehicle control, n=11 for indisulam, n=12 for olaparib, and n=10 for drug combination) were monitored until end of study (day 120) or when humane endpoint (as per project license) was reached. At endpoint, omental tumor deposits were collected and snap-frozen in liquid nitrogen and stored at -80°C until further analysis. Five mice survived to day 120. In four of the five mice, we could not guarantee successful tumour implantation (such as developing subcutaneous tumour deposits after injection of ID8 cells) and were excluded from the Kaplan-Meier survival analysis.

### Statistical analysis

Statistical analyses were performed by GraphPad Prism 9, except proteomics analysis (Perseus 1.6.15.0). For data with normal distribution, the Student’s t-test was used for statistical analysis between two groups. One-way ANOVA and multiple comparisons test were used for analysis among three or more groups. Two-way ANOVA and multiple comparisons test were used when there were three or more groups with two factors (for example, when there were more than two treatments with multiple concentrations). For skewed data, Kruskal-Wallis test and multiple comparisons test were used. The Log-rank (Mantel-Cox) test with Bonferroni corrected significance threshold (0.0083) was used to calculate statistical differences between surviving proportions in the animal studies.

## Acknowledgments

We thank animal technicians from the Imperial College London CBS animal center and Fabio Grundland-Freile for supporting animal studies. We thank Imperial NIHR BRC Genomics Center for conducting the RNA sequencing.

## Author contributions

Conceptualization: YuX, IM, HCK, AN. Methodology: YuX, SS, IM, HCK, AN. Investigation: YuX, SS, YM, MPL, MG, FM, YiX, CW, MRS, AN. Visualization: YuX, MPL, AN. Funding acquisition: HCK, AN. Project administration: HCK, AN. Supervision: IM, HCK, AN. Writing original draft: YuX, HCK, AN. All authors contributed to the review and editing of the paper.

## Data and materials availability

The *DCAF15* copy number variation and mRNA expression data in serous ovarian cancer patients were generated by the TCGA Research Network (PanCancer Atlas) and downloaded from the cBioPortal (https://www.cbioportal.org). The CTD^2^ Drug sensitivity data in ovarian cancer cell lines and *DCAF15* expression data were downloaded from the DepMap (https://depmap.org/portal/). The alternative splicing data included for systematic literature analysis were downloaded from online versions of the original publications. All other data are available in the main text or the supplementary materials.

