## Supplementary file for "Pharmacological depletion of RNA splicing factor RBM39 by indisulam synergizes with PARP inhibitors in high-grade serous ovarian carcinoma"

**Affiliations:**

### SUPPLEMENTARY MATERIALS

### MATERIALS AND METHODS

#### Materials

Indisulam (SML1225) and E7820 (SML2950) were purchased from Sigma-Aldrich and dissolved in dimethyl sulfoxide (DMSO) for *in vitro* use. Olaparib (S1060), rucaparib phosphate (S1098), veliparib (S1004), and niraparib (S2741) were supplied by Selleckchem and reconstituted in

DMSO for *in vitro* use. The IdU (I7125) and BrdU (B5002) were purchased from Sigma-Aldrich and dissolved in 1M $NH_4OH$ and DMSO, respectively. Z-VAD-FMK (APExBIO, #A1902) was dissolved in DMSO. All drugs were prepared and stored according to manufacturers' instructions.

**Cell lines and cell culture**

KURAMOCHI was acquired from Dr. Haonan Lu, Imperial College London. OVSAHO and OVISE were purchased from the Japanese Collection of Research Bioresources (JCRB) cell bank. COV318, COV504, and COV644 were obtained from the European Collection of Authenticated Cell Culture (ECACC). The PEO1 cells for *DCAF15* knockdown or knockout experiments had a missense reversion mutation 5192A>T in *BRCA2*, and it was a kind gift from Professor Bob Brown's laboratory, Imperial College London. An independent stock of PEO1 (without the reversion mutation) was purchased from the Public Health England and used for dose-response curves and synergy analysis without CRISPR-Cas9 editing. PEO4 was purchased from Public Health England. KURAMOCHI, COV318, COV504, COV644, and the parental PEO1 were authenticated by ECACC Public Health England (case #11232). OVSAHO, OVISE, PEO1 (purchased from Public Health England) and PEO4 were authenticated by the supplier. Mouse ovarian cancer cell line ID8-*Trp53*$^{-/-}$ and ID8-*Trp53*$^{-/-}$;*Brca2*$^{-/-}$ lines were directly obtained from Professor Iain McNeish's laboratory (*1*). All cell lines were tested regularly for mycoplasma contamination.

OVSAHO and OVISE cells were cultured in RPMI-1640 media (Gibco$^{TM}$) supplemented with 10% fetal bovine serum (FBS, Gibco$^{TM}$, or FirstLink), 2 mM of L-glutamine (Gibco$^{TM}$), and 100 units/mL of penicillin-streptomycin (Gibco$^{TM}$). PEO1-derived cells and PEO4 were cultured in RPMI-1640 supplemented with 10% FBS, 2 mM L-glutamine, 100 units/mL of penicillin-streptomycin, and 2 mM sodium pyruvate (Gibco™). COV318, COV504, COV644, and KURAMOCHI cells were cultured in DMEM media (Gibco™, without glucose or phenol red) supplemented with 10% FBS, 2 mM L-glutamine, 100 units/mL of penicillin-streptomycin, and 5.6 mM glucose (Gibco™). ID8 clones were cultured in DMEM media (without glucose or phenol red) supplemented with 4% FBS, 2 mM L-glutamine, 100 units/mL of penicillin-streptomycin, 25 mM glucose, and 1x Insulin-Transferrin-Selenium solution (Gibco™) as previously described (*1*). All cells were cultured at 37°C with 5% $CO_2$.

**CRISPR-Cas9 knockout of DCAF15**

PEO1 cells were seeded in 96-well plates and transduced with Edit-R CRISPR-Cas9-TurboGFP lentiviral particles (Dharmacon #VCAS11864). Transduced cells with GFP expression were sorted via FACS. PEO1-Cas9 cells were then expanded and transfected with crRNAs and Edit-R CRISPR-Cas9 synthetic tracrRNA (Dharmacon #U-002005) using lipofectamine 2000 (Life Technologies). To improve gene editing efficiency, a cocktail of three crRNAs targeting *DCAF15* was used, consisting of (1) 5′-GGAGACCCAGAAGAACGGGC-3′, (2) 5′-GCAGCTTCCGGAAGAGGCGA-3′, (3) 5′- ACAGCAAGCTCAAGCTG-3′. Edit-R crRNA Non-Targeting Control (Dharmacon #U-007501-01) was used alongside as a blank control. Following transfection, single-cell clones were expanded and sequenced for *DCAF15* editing. DCAF15 knockout was also validated using indisulam dose response studies. Loss of genomic DCAF15 was confirmed by sequencing.

**Systematic literature analysis**

A PubMed search using the keyword "RBM39" was performed on 10 August 2022. Inclusion criteria were defined as research articles conducted RNA sequencing in RBM39-depleted human cancer cell lines, and the list of alternatively spliced genes is publicly available. Exclusion criteria were non-research articles, non-human cancer cell lines, and non-publicly available alternative splicing data. After full-text reading, five articles containing 11 lists of alternatively spliced genes were included for analysis (*2-6*). Exon-skipped genes and intron-retained genes were extracted from each list. The occurrence of unique genes (exon-skipped or intron-retained) in was counted across 11 lists. The common splicing targets of RBM39 were defined as exon-skipped genes occurred at least six times, or intron-retained genes occurred at least six times. Duplicates were removed for enrichment analysis.

**RNA sequencing and data analysis**

KURAMOCHI cells were treated with 0.1% DMSO or 10 μM indisulam for 24 hours. The total RNA was extracted using the miRNeasy Mini Kit (Qiagen, #217084) following manufacturer's protocol. RNA samples of three biological repeats were sent for sequencing at the National Institute for Health and Care Research-funded Imperial Biomedical Research Centre (BRC) Genomics Facility. All samples passed Quality Control assessment using a Tapestation. Samples were sequenced using a Nextseq2000 P2 (200 cycles) sequencer, and 100 bp paired-end reads were generated. Quality control checks were performed using FastQC. The GRCh38 reference genome (unmasked primary assembly) and GTF file (release 107) were downloaded from Ensembl (http://www.ensembl.org/info/data/ftp/index.html). STAR was used to align reads to the reference genome and read counts per gene were determined using featureCounts. The alternative splicing analysis was performed SpliceFisher (*7*) and rMATS algorithms (*8*). Sashimi plots were generated using Integrative Genomics Viewer (IGV, version 2.15.2, Broad Institute). Exon annotations were based on the GRCh38 human reference genome (http://uswest.ensembl.org/index.html). RNAseq data has been uploaded to the NCBI Gene Expression Omnibus and are accessible through GEO Series accession number GSE223011.

**Functional annotation and enrichment analysis**

The Database for Annotation, Visualization and Integrated Discovery (DAVID, https://david.ncifcrf.gov/summary.jsp) (*9, 10*) was used for functional annotation and enrichment analysis with four databases selected: Gene ontology biological processes (DIRECT), Kyoto Encyclopedia of Genes and Genomes (KEGG), REACTOME, and WIKIPATHWAY. A query-specific background gene list was used whenever possible. Significant enrichment was defined as terms or pathways with FDR less than 0.1.

**PCR-based alternative splicing analysis**

Total RNA from cells was extracted using the RNeasy Mini Kit (Qiagen, #74106) or miRNeasy Mini Kit (Qiagen, #217084) following the supplier's instructions with on-column DNase digestion. Equal quantities of RNA were reverse transcribed to cDNA using the High-Capacity RNA-to-cDNA kit (Applied Biosystems). To detect alternative splicing variants, we designed primers flanking the regions of interest (sequence provided below) and amplified them using the Q5 High-Fidelity PCR kit (New England Biolab). PCR reactions included 35 cycles, and annealing temperatures were calculated using the annealing temperature calculator (https://tmcalculator.neb.com/#!/main). PCR products were electrophoresed using 2-3% agarose

(Sigma-Aldrich) gels mixed with 1:10,000 SYBRsafe DNA gel stain (Invitrogen). Gel images were visualized with a GelDoc.

Primers used in this study included *ATM* (exon 51 to 54) forward 5'-TGCCTCTTATGTACCAATTGGCT-3', reverse 5'-AATTGGCTGGTCTGCTGGAA-3'; *BRCA1* (exon 7 to 10) forward 5'-AACTCTGAGGACAAAGCAGC-3', reverse 5'-CTGTAATGAGCTGGCATGAGT-3'; *RAD51D* (exon 3 to 6) forward 5'-TGGCTCAGTTCTCGGCTTT-3', reverse 5'- AGATGTCAAATGCATGCACCA-3'; *TP53BP1* (exon 15 to 18) forward 5'- GCTGTTGCTGAGTCTGTTGC-3', reverse 5'-TGCGTACTTCCCGGATTGTT-3'; *RIF1* (exon 21 to 23) forward 5'-AGGTGGGCAAAACTGGTCAT -3', reverse 5'-TGGGTGCTCCACTACGAAAT-3'; *FANCI* (exon 31 to 35) forward 5'- GAGTCAGGCCGAGAAGGTTC-3', reverse 5'-CAACGGCAGCAGGTTTCTC-3'; and *GAPDH* forward 5'-GGCTGCTTTTAACTCTGG-3', reverse 5'-GGAGGGATCTCGCTCC-3'.

**Quantitative Real-time PCR**

Total RNA from cell lines was extracted directly from culture flasks using the RNeasy Mini Kit (Qiagen, #74106) following the supplier's instructions. Equal quantities of RNA were reverse transcribed to cDNA using the High-Capacity RNA-to-cDNA kit (Applied Biosystems). DCAF15 gene expression was measured using TaqMan gene expression probes (Hs00384913_m1) normalized to BACT (Hs01060665_g1) according to manufacturer's guidelines (ThermoFisher). Analysis was conducted using the comparative CT method (*11*).

**Protein extraction**

Total proteins of cultured cells were extracted using RIPA lysis buffer (Sigma-Aldrich, #R0278) supplemented with at least 1% HALT protease and phosphatase inhibitor cocktail (ThermoFisher, #78442). Proteins from tumor tissues were extracted in RIPA buffer supplemented with 3% protease and phosphatase inhibitor cocktail at a rate of 50-100μL lysis buffer per 10 mg tissue. To facilitate protein extraction, tissues were mixed with glass grinding beads and homogenized by the Cryolys Evolution homogenizer, followed by sonication in an ice bath for 10 – 20 minutes. Protein supernatants were collected after centrifugation at 20,000xg at 4°C for 20 minutes and stored at -80°C until analysis.

**Western blot**

An equal amount of protein was loaded onto triglycine Mini-PROTEAN precast gels (Bio-Rad) for electrophoresis (200V for 35 to 50 minutes), followed by wet transfer (100V for 60 minutes) to nitrocellulose membrane via wet transfer. Membranes were blocked in 5% skimmed milk in tris-buffered saline containing 0.1% of Tween-20 (TBST) or 5% bovine serum albumin (in TBST, for phosphorylated targets) for 60 minutes at room temperature. Primary antibodies were diluted in 5% skimmed milk and incubated overnight at 4 °C. Horseradish peroxidase (HRP)-conjugated secondary antibodies were diluted in 5% skimmed milk and incubated for 60 minutes at room temperature. Before and after each antibody incubation, membranes were washed three times with TBST. Proteins of interest were detected with the SuperSignal West Pico PLUS chemiluminescent substrate (Thermo Scientific) and Amersham Hyperfilm ECL (GE Healthcare). Image J Fiji V2.0.0 was used for protein bands densitometry analysis. Membranes might be stripped once with the Restore WB stripping buffer (Thermo Scientific, #21059) and re-probed with a different pair of primary and secondary antibodies.

Primary antibodies used in this studies were rabbit anti-RBM39 (1:500, Sigma-Aldrich, #HPA001591), mouse-anti-β-actin (1:10,000 - 1:50,000, Abcam, #ab6276), mouse anti-phospho-Histone H2A.X (ser139) γH2AX (1:1000, Sigma-Aldrich, #05-636), rabbit anti-human pChk2 (T68) (1:300, R&D system, AF1626), rabbit anti-human Chk2 (1:500, R&D system, MAB1358), rabbit anti-pATM-S1981 (1:2000, Abcam, #ab81292), mouse anti-ATM (1:1000, GenTex, GTX70103), rabbit anti-pChk1-S345 (Cell Signaling Technology, CST2348, 1:1000), and mouse anti-Chk1 (Cell Signaling Technology, CST2360). HRP-conjugated secondary antibodies included goat anti-rabbit polyclonal antibody (1:2500 - 1:5000, Invitrogen, #31460), and goat anti-mouse polyclonal antibody (1:2500 – 1:5000, Invitrogen, #31340).

**Sulforhodamine B (SRB) cell growth assay**

SRB assay was conducted as previously reported (*12*). SRB dye was supplied by Sigma-Aldrich. For a 48 or 72-hour growth assay, cells were counted using a hemocytometer (Hawksley) and seeded in 96-well plates at the following densities: 500 cells per well (ID8 cells), 2000 cells per well (OVISE, COV318, and COV504), 2500 cells per well (PEO1-derived cells), 4000 cells per well (PEO4, COV644, and KURAMOCHI), and 5000 per well (OVSAHO). Cells were allowed to attach overnight before drug treatment the next day. At the end of drug treatment, cells were fixed with 10% (w/v) of trichloroacetic acid (TCA) solution at 4°C for 60 minutes and rinsed gently under tap water. Plates were left on the bench to dry before staining with 0.4% SRB solution (in 1% acetic acid) for 30 minutes. Excess SRB dye was washed away with 1% acetic acid. After drying on the bench, 10 mM Tris-base was added to each well to solubilize the protein-bound dye. Cell growth was measured using absorbance at 565 nm or 510 nm with a plate reader (OPTImax Molecular Devices or BMG Labtech ClarioStar). A blank control without cells was always included for background subtraction. Cell growth values under drug treatment were normalized to vehicle control (0.1% – 0.2% DMSO).

**Drug combination and synergy analysis**

Drug-drug interactions were assessed using the Chou-Talalay method (*13-15*). A fixed combination ratio for each drug pairs was experimentally determined by concentrations that caused similar cytotoxicity (30% to 50% growth inhibition). CI values were calculated using the CompuSyn (*15*). $CI < 1$ indicates synergism, $CI = 1$ indicates additive, and $CI > 1$ indicates antagonism.

**Caspase 3/7 activation apoptosis assay**

Caspase-Glo 3/7 assay (Promega) was used to measure caspase 3/7 activities following the manufacturer's instructions. Cells were seeded and treated in the same way as SRB assays. After 48-hour treatment, the conditioned media was aspirated and replaced with 30-50 µL of fresh media. An equal volume of the Caspase-Glo 3/7 reagent was added and incubated on a shaker for at least 45 minutes. The reaction mixture from each well was then transferred to an opaque-walled 96-well plate for measurement by a ClarioStar plate reader (BMG Labtech). A cell-free control was involved for background correction. A parallel plate was always included to quantify cell mass by SRB assay, which was used to normalize caspase 3/7 activities.

**Annexin V cell apoptosis analysis**

After drug treatment, adhering cells were detached from plates by trypsinization (TrypLE Express, Gibco, #12604013) and combined with floating cells from the conditioned media. After two washes with cold phosphate-buffered saline (PBS), cells were resuspended in Annexin V binding buffer (0.01M Hepes, 0.14M NaCl, 2.5 mM $CaCl_2$) and passed through a 30 µm cell strainer to generate single-cell suspensions. A staining solution containing Cy5.5-conjugated Annexin V (2.5µL per test, BD Biosciences, #559925) and PI solution (2.5 µL per test, Invitrogen, #00-6990-50) was used to stain cells. After a 15-minute incubation in the dark, stained cells were analyzed with a BD FACS Canto analyzer. Positive controls for each staining were included in each repeat for color compensation. FlowJo V10.6.2 was used for gating and analysis. The same gating was used for biological repeats.

**Colony formation assay**

Suspensions of OVSAHO cells were passed through a 40 µm cell strainer (Corning) to generate single-cell suspensions and seeded in 6-well plates at a density of 3000 cells per well. Cells were allowed to grow for five days to form small colonies before drug treatment for 48 or 72 hours. Immediately after drug treatment, cells were washed twice with PBS and incubated in fresh media. The culture media was refreshed every 2-3 days until 16 days post-seeding. On day 16, colonies were fixed in ice-cold methanol for 10 minutes and stained with 0.5% crystal violet (in 20% methanol) for another 10 minutes. Plates with fixed colonies were rinsed gently under tap water and left on the bench to dry overnight. Plates were imaged using a scanner, and the Fiji V2.0.0 was used to count colonies. Each well was manually cropped out from the entire plate and converted into 8-bit images. Colony numbers were measured following these steps: (1) highlighting edges of colonies by “Find edge”; (2) adjusting the color threshold to 42 – 255; (3) colonies were automatically counted by the “Analyze Particle” function using these parameters: particle size >

50 pixels (in a 1.35 inches x 1.35 inches, 800 pixel/inch dimension image) and circularity between 0.3 and 1.0.

### Cell cycle analysis

In the last 60 minutes of drug treatment, cells were incubated with 10 μM BrdU and 200 μM IdU sequentially for 30 minutes each. Three washes with lukewarm PBS were included between BrdU and IdU labeling. This labeling strategy was selected because it generates cells suitable for both cell cycle and DNA fiber analysis. After labeling, cells were detached by trypsinization and washed twice with PBS. To fix cells, freshly prepared 70% ethanol was added dropwise to the cell suspension and incubated at 4°C overnight. After removing ethanol, 2M HCl with 0.5% Tx-100 was added dropwise to the cell pellet and incubated at room temperature for 30 minutes with agitation to denature chromosomes. Then, cells were washed with 0.1M $NaB_4O_7$ (pH 8.5) for five minutes to neutralize residue HCl, followed by blocking in the blocking buffer (1% BSA in PBS with 0.2% Tween-20) for 30 minutes. Incorporated BrdU and IdU were probed by mouse anti-BrdU (1:25 in blocking buffer, BD347580, cross-reacts with IdU) and AF488-conjugated goat-anti-mouse antibody (1:100 in blocking buffer, Invitrogen, #A11001) for 60 minutes each at room temperature. Two washes with the blocking buffer were included between primary and secondary antibody incubations. Afterward, cells were incubated with FxCycle PI/RNase staining solution (Invitrogen, #F10797) for 15 minutes in the dark. Finally, stained cells were analyzed on a BD FACS Canto analyzer, and data were analyzed using Flowjo V10.6.2. Color compensation was performed for every experiment, and the same gating were used for biological repeats.

### DNA fiber analysis

DNA fiber analysis was performed as previously reported (*16, 17*). Cells were pulse-labeled with 10 μM BrdU and 200 mM IdU in the same way as the cell cycle analysis. Three washes of lukewarm PBS were included between pulse labeling. Cell suspensions were washed with ice-cold PBS and kept in frozen media at -80°C until further analysis. On the day of DNA fiber analysis, cells were thawed at 37°C and washed with ice-cold PBS to remove the freezing media. Approximately 2000 cells were spotted on a SuperFrost slide (2000/2 μL/spot, 2 spots/slide). The drop was allowed to dry slightly for five minutes and immediately mixed with 7 μL of lysis buffer (200 mM Tris-HCl, pH 7.5, 50 mM EDTA, and 0.5% SDS) to release DNA from cells. After airdrying for five to seven minutes, slides were tilted at about 15°-30°, enabling the DNA mixture to run down the slide slowly. Next, slides were dried for 30 minutes, followed by 10 minutes of fixation (in 75% methanol and 25% acetic acid) and drying for another 30 minutes. Dried DNA spreads were briefly rinsed with distilled water and immersed in 2.5M HCl for one hour to denature DNA. After two PBS washes, slides were blocked in the blocking buffer (1% BSA in PBS with 0.1% Tween-20) for one hour. BrdU was probed first by incubating slides with rat anti-BrdU antibody (1:50, Abcam, #ab6326) for 90 minutes, followed by two PBS washes, one wash with blocking buffer, and then one-hour incubation with AF647-conjugated goat anti-rat antibody (1:100, Invitrogen, #A21247) in the dark. Slides were washed and left in PBS overnight before IdU labeling. The next day, IdU was detected by one-hour incubation of mouse anti-BrdU antibody (for IdU detection, 1:50, #BD347580), three washes as above, and one-hour incubation of AF488-conjugated goat anti-mouse antibody (1:100, Invitrogen, #A11001). After rinsing away excess antibodies, slides were mounted with Fluoromount-G (Invitrogen, #00-4958-02), covered with a coverslip, and sealed with nail polish. Slides were imaged using the Aurox laser-free confocal microscope within 48 hours. Positions of each imaging field were recorded to avoid duplicating fibers being captured. DNA track length was manually measured using Fiji V2.0.0. All antibodies were diluted in the blocking buffer.

**Immunofluorescence analysis**

Cells were seeded and treated in 96-well plates. At the end of the drug treatment, cells were washed once with PBS and fixed with 4% formaldehyde (Fisher Scientific, #11586711) for 15 minutes. After removing formaldehyde and two PBS washes, cells were permeabilized with 0.1% Triton-X-100 (Sigma-Aldrich, #T8787) in PBS for 20 minutes. Permeabilized cells were washed with PBS once before blocking in the blocking buffer (1% BSA and 2% FBS in PBS) for 60 minutes. Cells were incubated with mouse anti-γH2AX antibody (1:500 in blocking buffer, Sigma-Aldrich, #05-636) overnight at 4°C, followed by three PBS washes on a rocker. Next, cells were incubated in staining solutions containing AF488-conjugated goat anti-mouse antibody (1:500 in blocking buffer, Invitrogen, #A11001) and DAPI (1:1000 in blocking buffer, Sigma-Aldrich, #D9542) for two hours at room temperature. Stained cells were washed with PBS three times and 200 µL and imaged in 200 µL PBS. IN Cell Analyzer 1000 (Cytivia, UK) was used for imaging using a 20X objective. A minimum of 16 fields per well were acquired and CellProfiler (V3.1.8.) was used for image analysis.

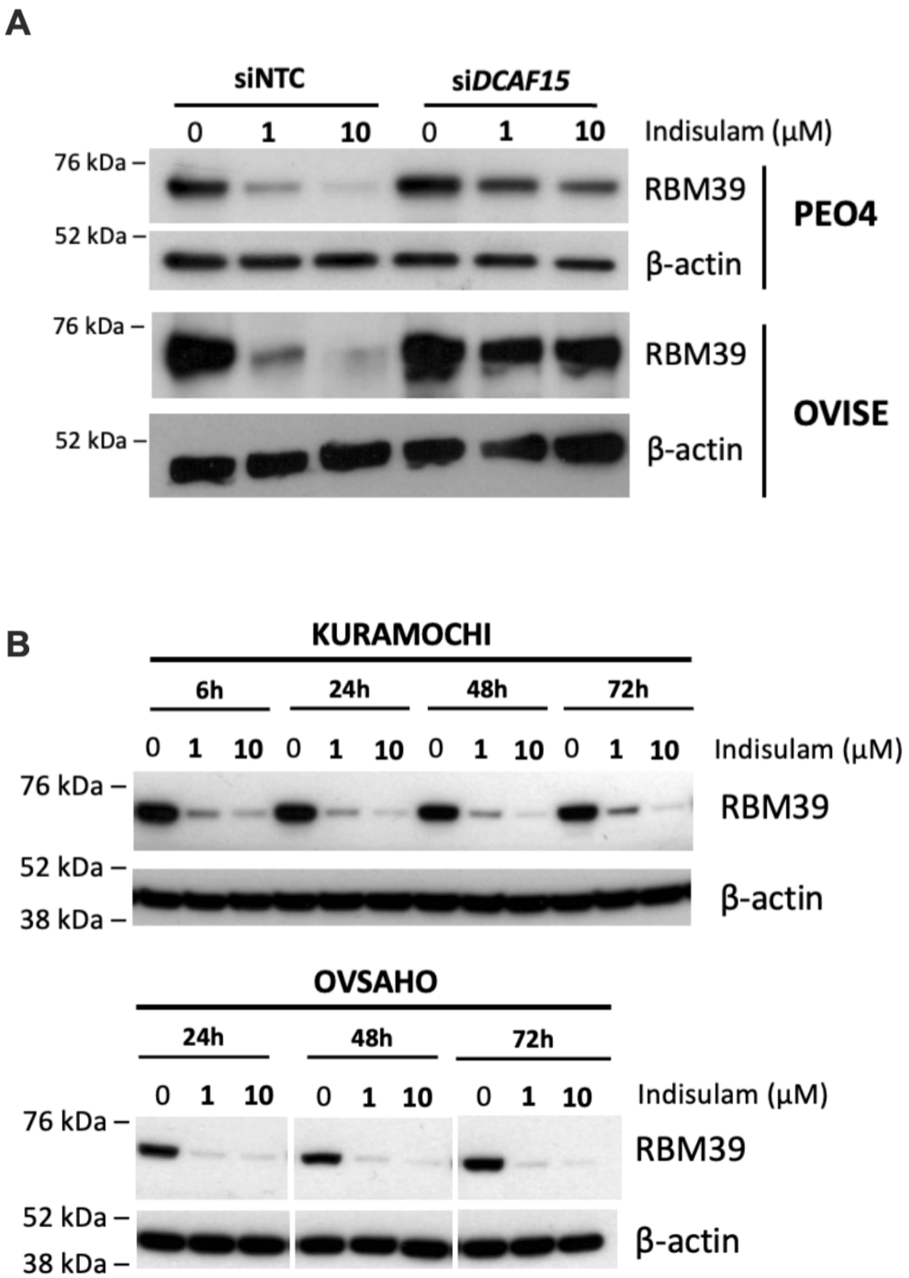


**Fig. S1. DCAF15 is essential for indisulam to degrade RBM39. (A)** *DCAF15* in PEO4 or OVISE cells were knocked down using siRNAs, and RBM39 levels were investigated using western blot (representative image of *n*=3 individual experiments). **(B)** KURAMOCHI (*n*=1) and OVSAHO (*n*=1) cells were treated with indicated concentrations of indisulam for 6 to 72 hours and RBM39 abundance was assessed using western blot. β-actin was included as a loading control.

A

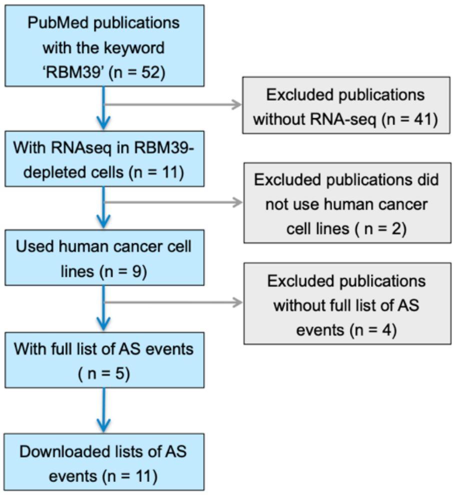


B

| No. | First author | Year | Disease | Cell line | Method of RBM39 depletion |
|---|---|---|---|---|---|
| 1 | Nijhuis, A. | 2022 | Neuroblastoma | IMR-32 | 5 μM Indisulam for 16 hours |
| 2 | Singh, S. | 2021 | Neuroblastoma | BE2C | shRBM39 (#188) for 72 hours |
| 3 | | | | BE2C | shRBM39 (#190) for 72 hours |
| 4 | | | | SKNAS | shRBM39 (#188) for 72 hours |
| 5 | | | | SKNAS | shRBM39 (#190) for 72 hours |
| 6 | Lu, S. X. | 2021 | Melanoma | 501-MEL-ES | 1 μM indisulam for 96 hours |
| 7 | | | | A375 | 1 μM indisulam for 96 hours |
| 8 | | | | SK-MEL-239 | 1 μM indisulam for 96 hours |
| 9 | Wang, E. | 2019 | Acute myeloid leukemia | MOLM-13 | CRISPR knockout, 6 days post transduction |
| 10 | | | | THP-1 | CRISPR knockout, 6 days post transduction |
| 11 | Mai, S. | 2016 | Breast cancer | MCF-7 | RBM39 knockdown for 48 hours |

C

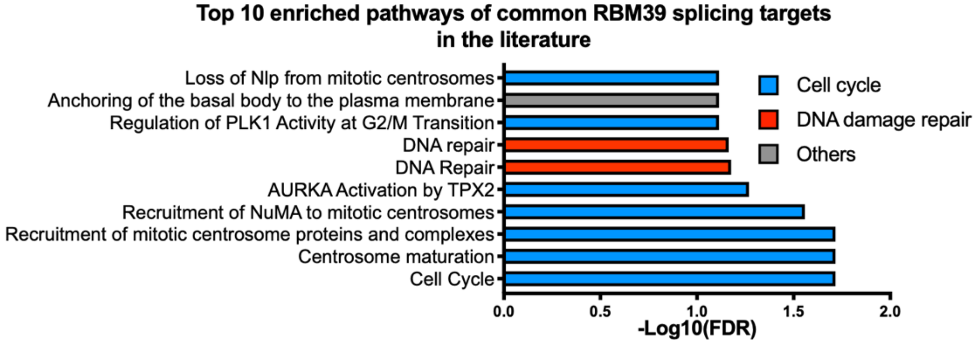


D

KURAMOCHI (SpliceFisher analysis)

| Alternative splicing events | Gene count |
|---|---|
| Intron retention | 8389 |
| Exon skipping | 7904 |
| Alternative 3' splice site | 841 |
| Alternative 5' splice site | 682 |
| Mutually exclusive exons | 319 |

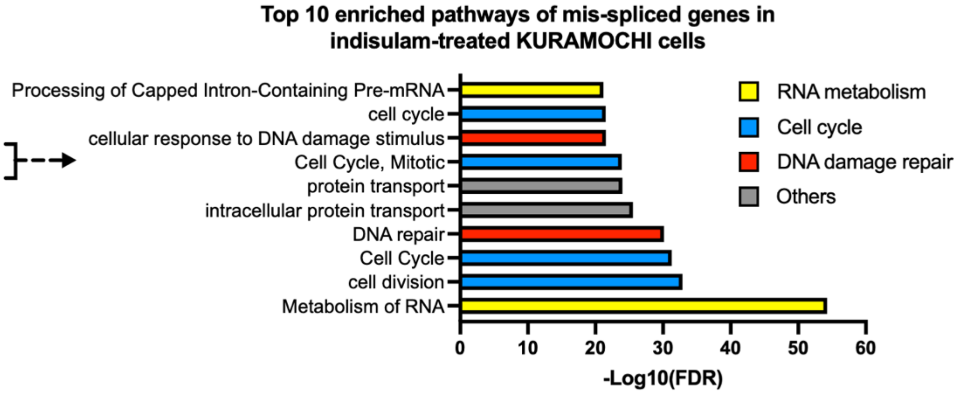


**Fig. S2. RBM39 regulates the splicing of cell cycle and DNA damage repair genes across cancer types. (A)** A flow chart showing the process of literature searches. **(B)** Key characteristics of eligible papers (*39, 44, 45, 51, 56*). **(C)** The unique common splicing targets of RBM39 (exon skipped or intron retained in at least six datasets) were used for pathway enrichment analysis. FDR

< 0.1 was considered significantly enriched. **(D)**. Alternative splicing events determined by SpliceFisher algorithm. Pathway enrichment analysis (FDR<0.1) on intron-retained or exon-skipped genes.

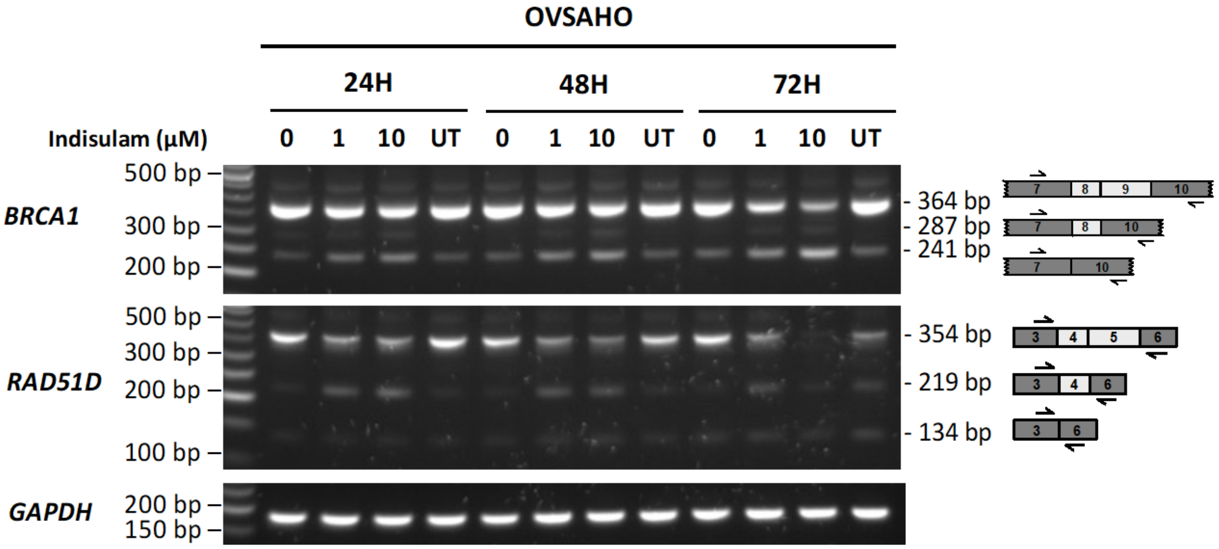


**Fig. S3. Indisulam induces mis-splicing of DNA repair genes in OVSAHO.** OVSAHO cells were untreated (UT) or treated with 0 µM, 1 µM or 10 µM of indisulam for 24 to 72 hours before RNA extraction. PCR-based alternative splicing analysis was performed to detect exon-skipping events of *BRCA1* and *RAD51D* (*n*=1).

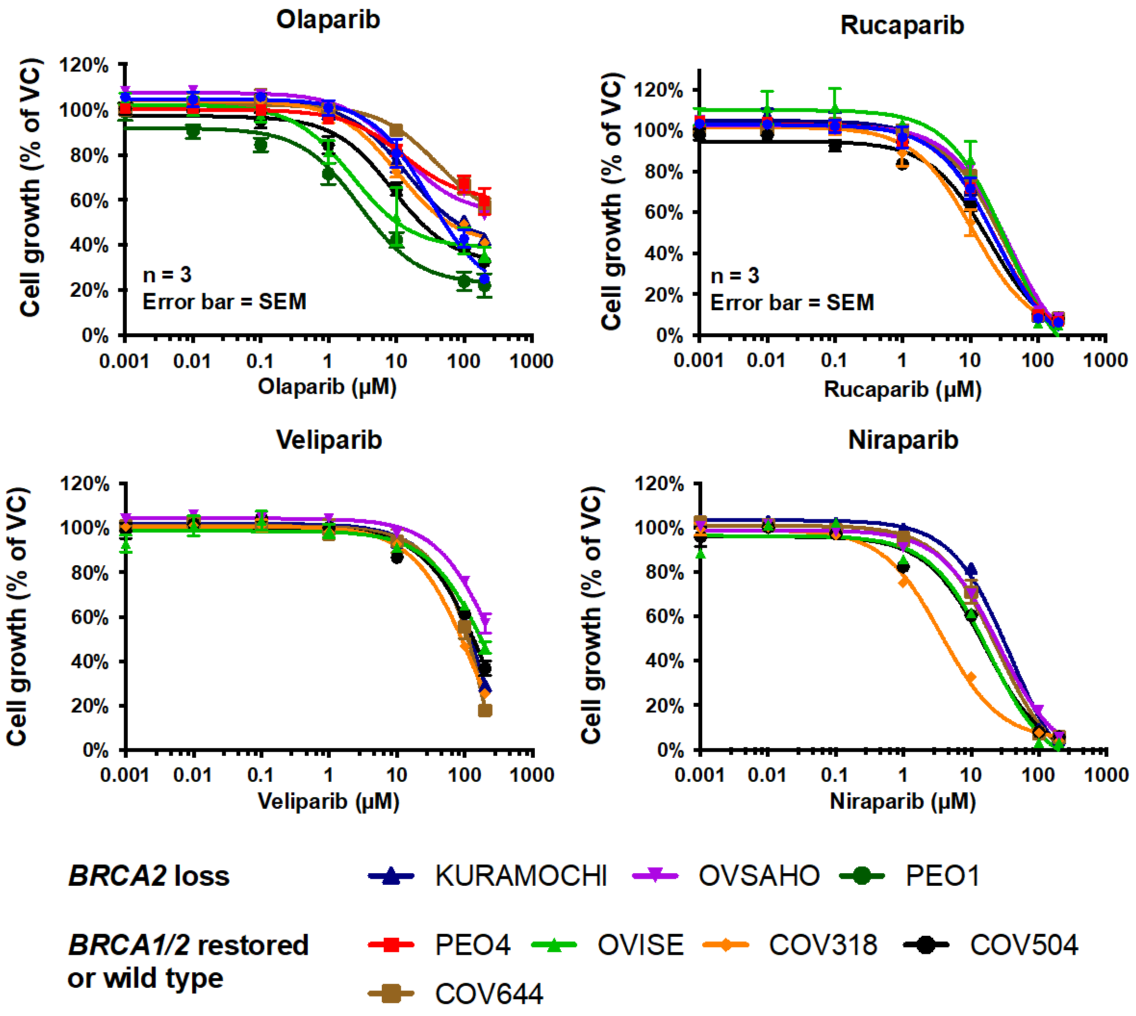


**Fig. S4. Sensitivity of human ovarian cancer cell lines to PARP inhibitors.** Dose-response curves of human ovarian cancer cell lines to four PARP inhibitors: olaparib, rucaparib, veliparib and niraparib. Data were collected from three independent repeats except veliparib dose-response curve for OVSAHO (*n*=2) and OVISE (*n*=2); niraparib dose-response for OVISE (*n*=1), OVSAHO (*n*=1) and COV318 (*n*=2). Error bar = SEM.

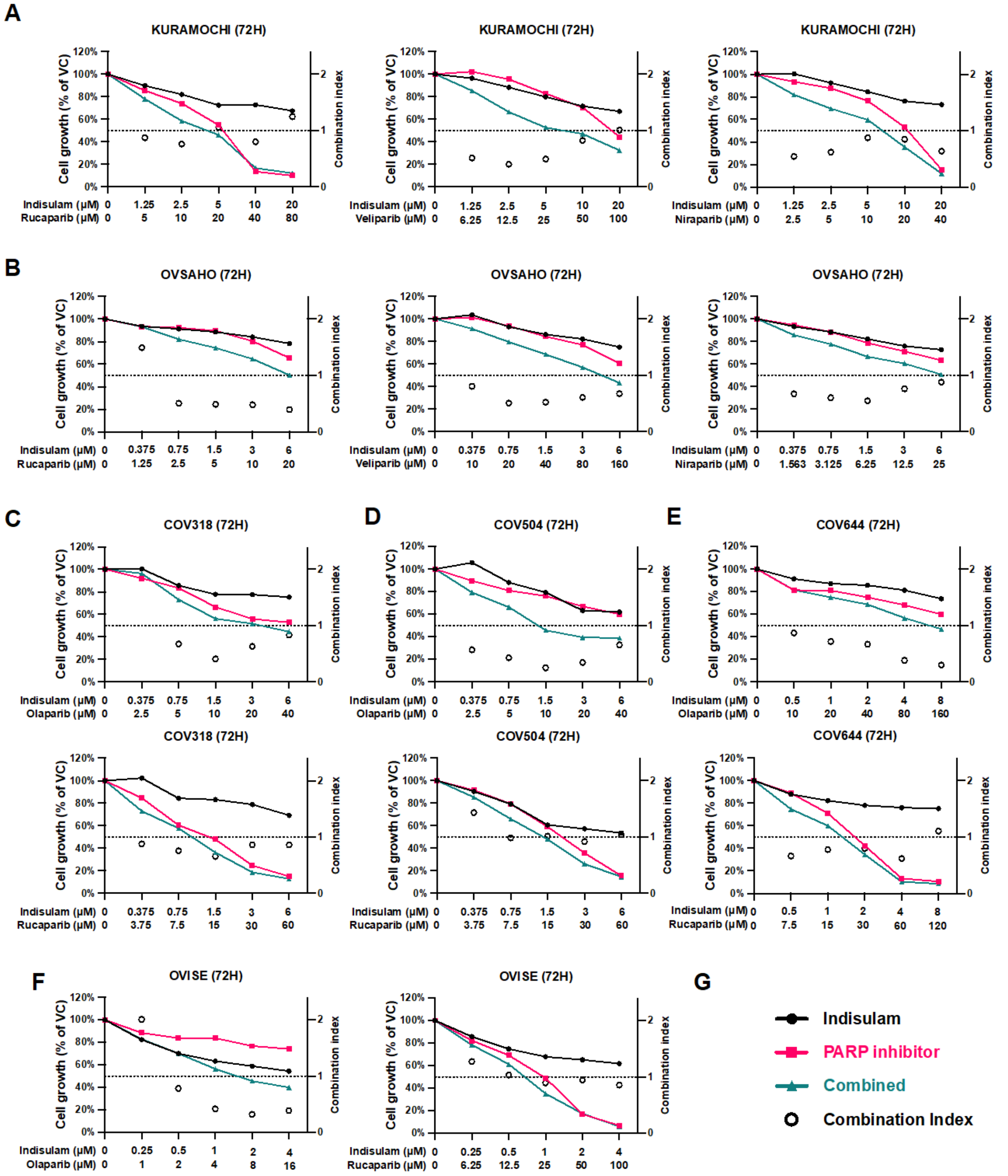


**Fig. S5. Indisulam synergizes with multiple PARP inhibitors in ovarian cancer. (A-B)** Cell growth of KURAMOCHI (a) and OVSAHO (b) treated with indicated doses of indisulam in combination with rucaparib, veliparib or niraparib. **(C-F)** Cell growth of COV318 (c), COV504

(d), COV644 (e) and OVISE (f) cells treated with indisulam in combination with olaparib or rucaparib. **(G)** The same color scheme was used for all graphs. The line graphs show the mean from three technical replicates to the left y-axis (*n*=1). Open circles indicate the CI values (to the right y-axis).

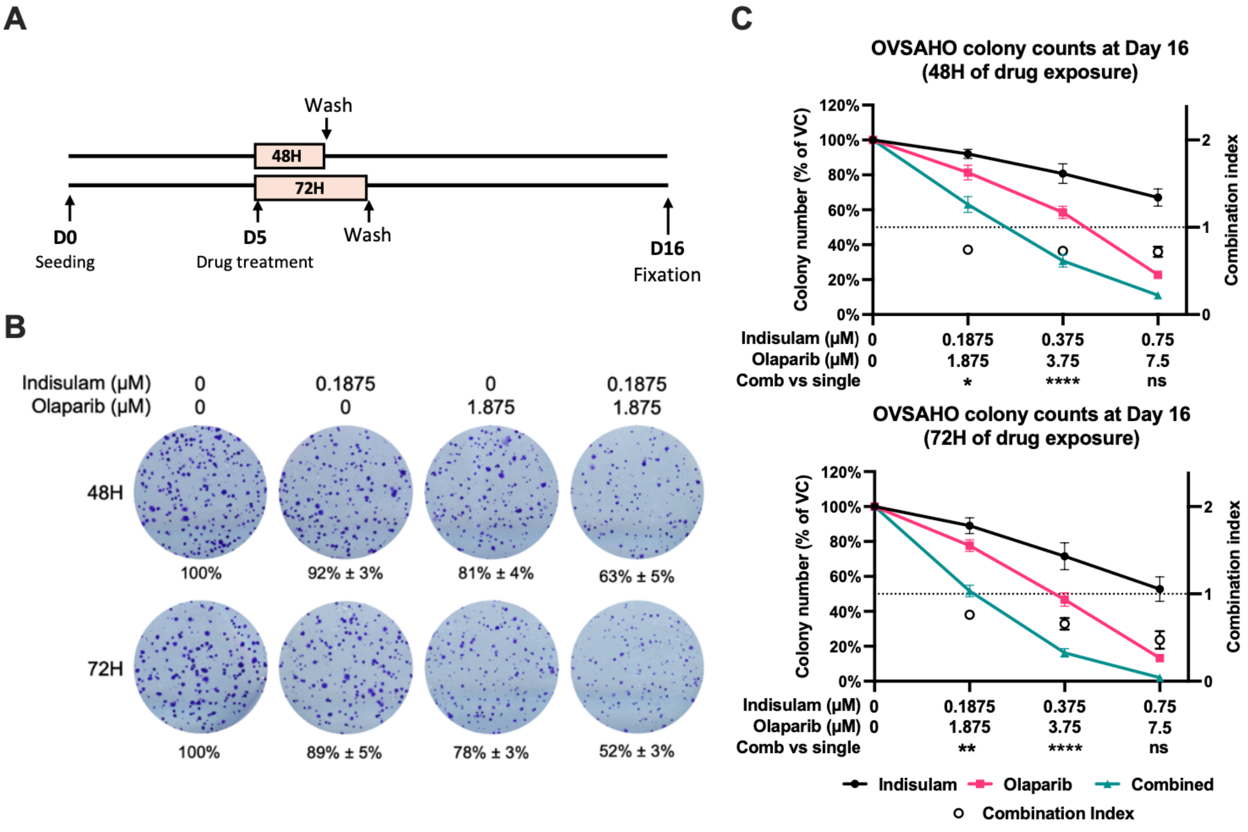


**Fig. S6. The combination of indisulam and olaparib inhibits OVSAHO clonogenicity. (A)** Schematic of the experimental design of the colony formation assay. OVSAHO cells were allowed to form colonies for five days before 48 or 72-hour exposure to indisulam, olaparib, or the combination, followed by further incubation in fresh media. Colonies were fixed on day 16 for counting. **(B)** Representative fields of OVSAHO colonies after fixation. **(C)** Measurements of colony number changes relative to vehicle control. At least three independent experiments were performed. The line graphs to the left y-axis indicate mean colony numbers relative to vehicle control (error bar = SEM), while open circles are mean CI values to the right y-axis (error bar =

SD). Two-way ANOVA and Dunnett's comparisons test were performed to determine the statistical significance between drug combination and monotherapies (ns, non-significant; *p< 0.05; **p<0.01; ***p<0.001; ****p<0.0001).

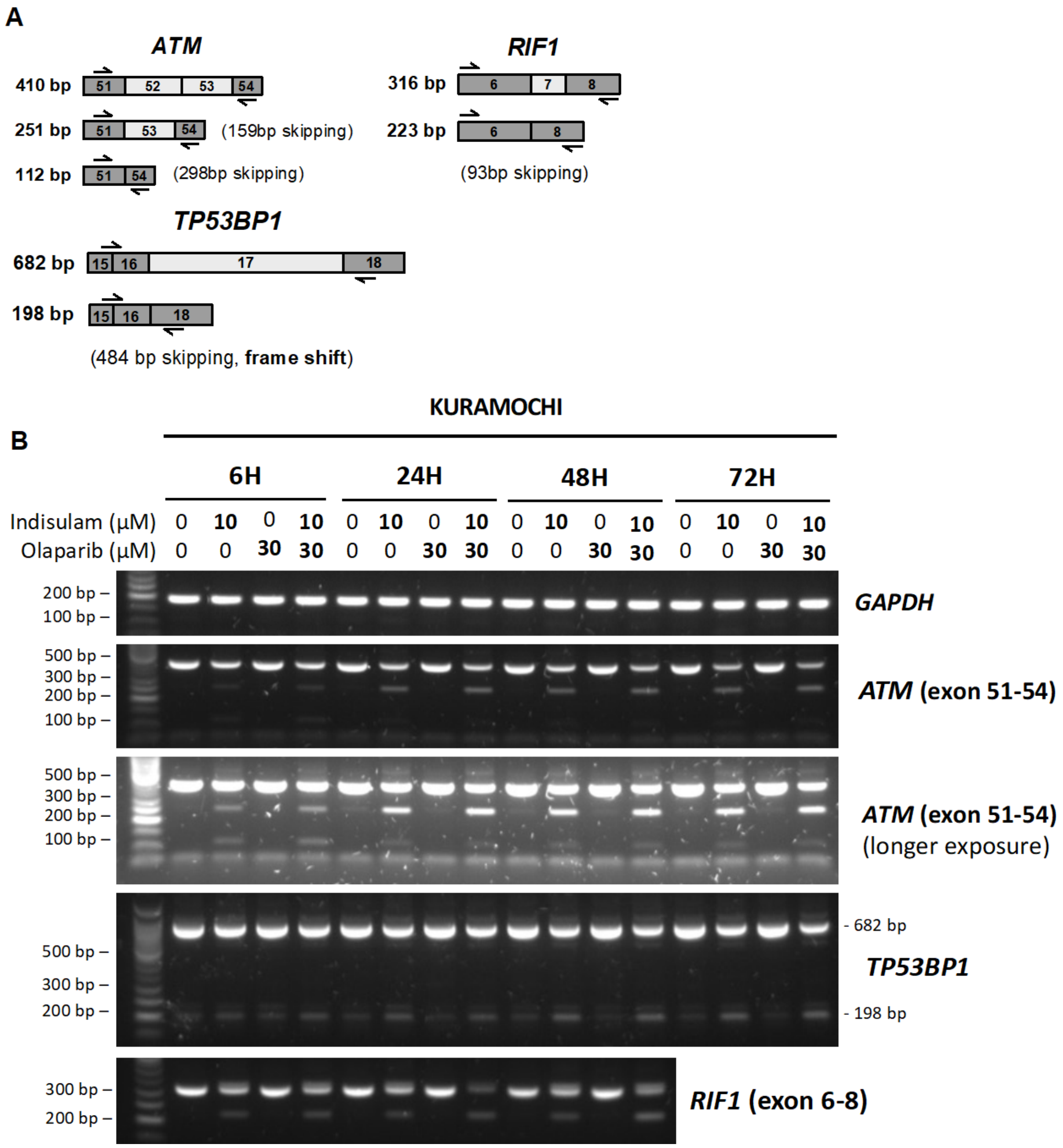


**Fig. S7. Indisulam-induced exon skipping of DNA damage repair genes are not affected in the presence of olaparib. (A)** Schematics showing primers designed to detect exon skipping events in *ATM, RIF1* and *TP53BP1*. **(B)** KURAMOCHI cells were treated with indicated drugs for 6 to 72 hours before PCR analysis for alternative splicing.

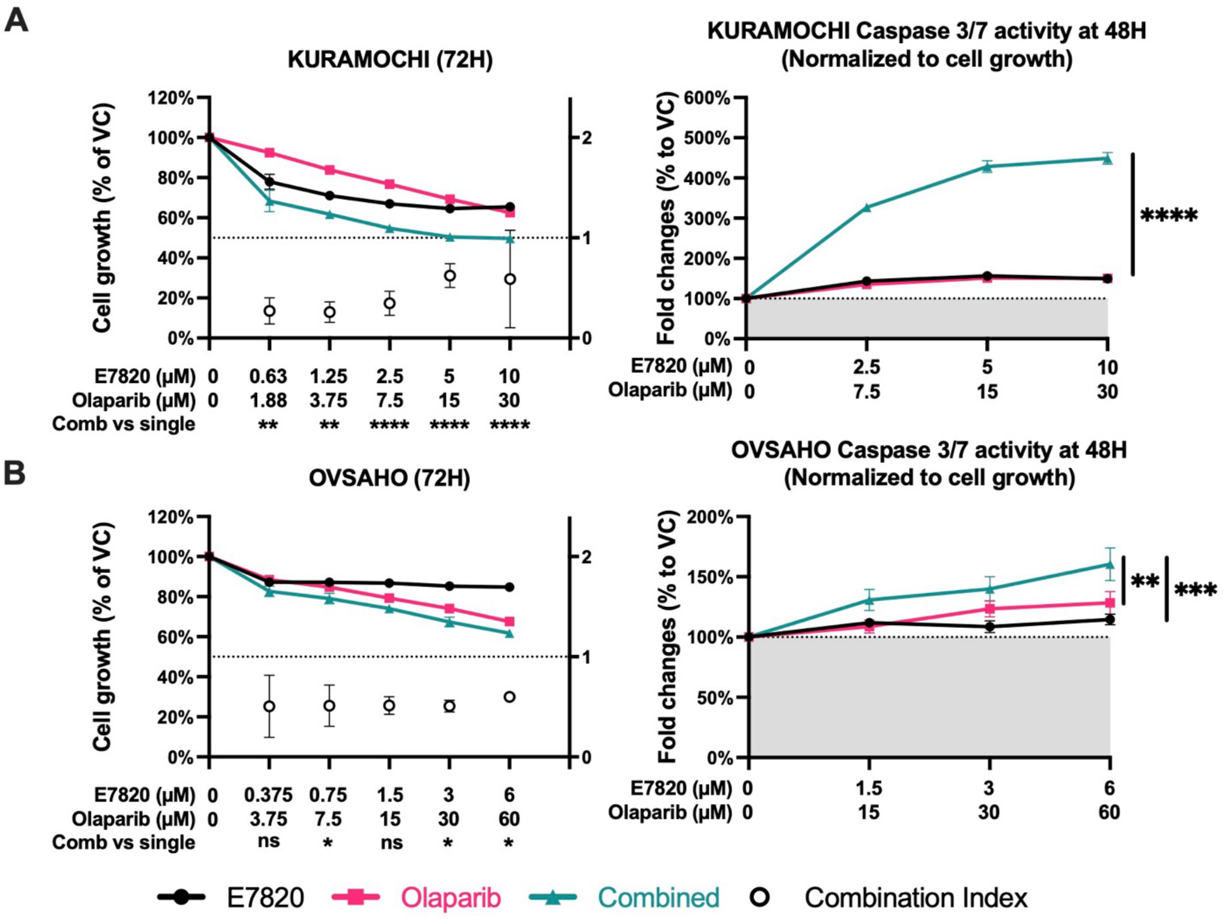


**Fig. S8. E7820 is synergistic with olaparib in HGSOC cells. (A)** KURAMOCHI cells were treated with indicated drugs. Cell growth was measured at 72 hours and cell apoptosis at 48 hours (*n*=3). **(B)** OVSAHO cells were treated with indicated drugs before measuring cell growth and apoptosis (*n*=3). Line graphs are to the left y-axis indicating mean cell growth relative to vehicle control (error bar = SEM), and open circles are to the right y-axis indicating mean CI values (error bar = SD). Grey areas indicate basal caspase activities in vehicle control. Two-way ANOVA analysis and Dunnett's comparisons test were performed to determine the statistical significance between the drug combination and monotherapies (ns, non-significant; *p<0.05; **p<0.01; ***p<0.001; ****p<0.0001).

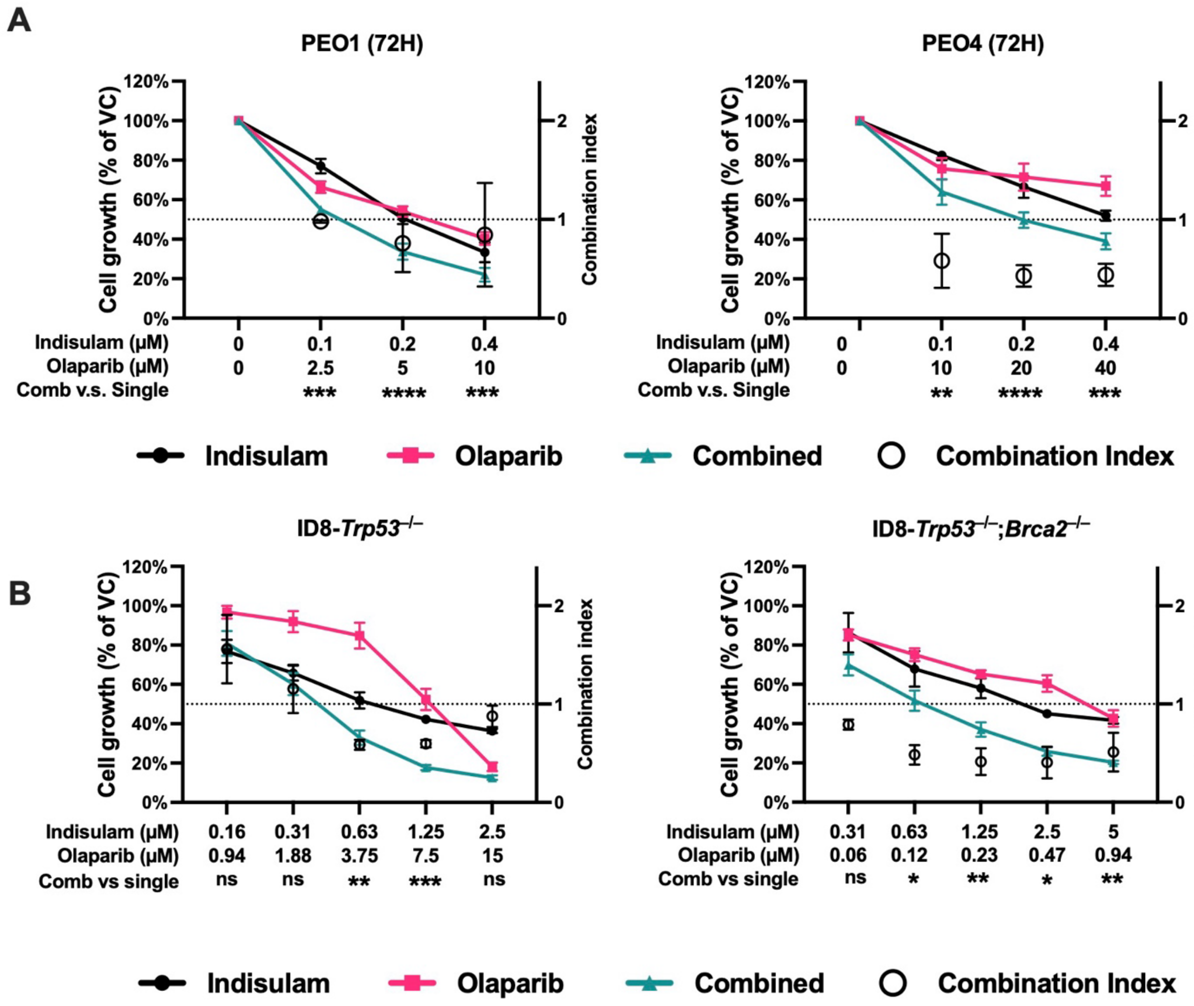


**Fig. S9. The synergy between indisulam and olaparib is effective regardless of *BRCA2* status.** **(A)** PEO1 and PEO4 cells were treated with indisulam and olaparib as monotherapies or in combination for 72 hours (n = 3). **(B)** ID8-*Trp53*$^{-/-}$ and ID8-*Trp53*$^{-/-}$;*Brca2*$^{-/-}$ clones were treated with indisulam and olaparib individually or in combination for 72 hours (*n*=3). Cell growth in (a) and (b) was measured with SRB assay. Line graphs are to the left y-axis indicating mean cell growth relative to vehicle control (error bar = SEM), and open circles are to the right y-axis indicating mean CI values (error bar = SD). The statistical significance between drug combination and monotherapies was calculated using two-way ANOVA and Dunnett's multiple comparisons test (ns, non-significant; *p<0.05; **p<0.01; ***p<0.001; ****p< 0.0001).

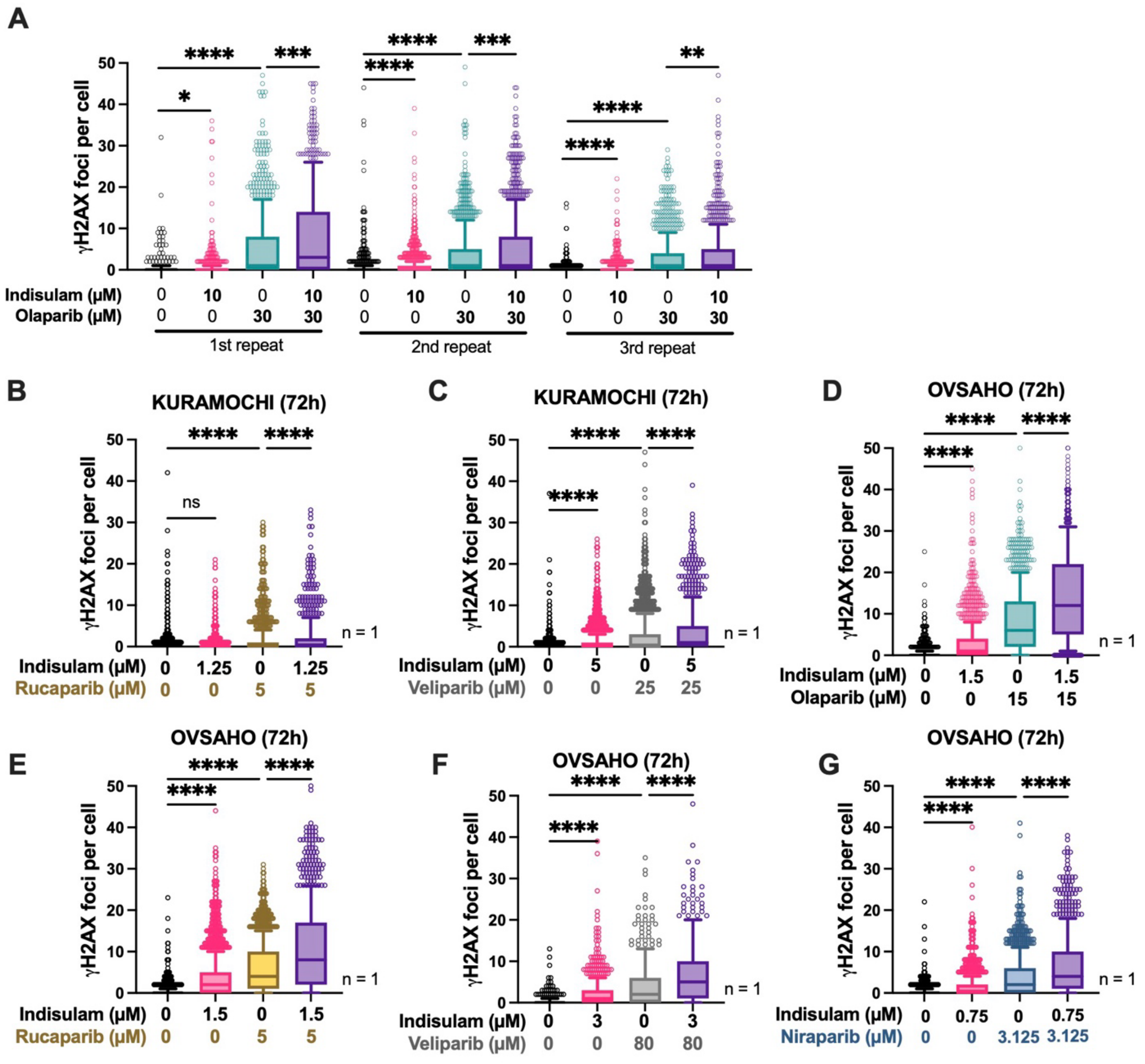


**Fig. S10. Indisulam increases γH2AX foci formation caused by PARP inhibitors. (A)** KURAMOCHI cells were treated with indicated drugs for 24 hours and γH2AX foci numbers in each cell were counted. Linked to Fig. 5B. **(B to G)** γH2AX foci counts in KURAMOCHI or OVSAHO cells after treatment of indisulam in combination with different PARP inhibitors (*n*=1). The Kruskal-Wallis test and Dunn's multiple comparisons test were used for statistical analysis (*p<0.05, **p<0.01, ***p<0.001, ****p<0.0001).

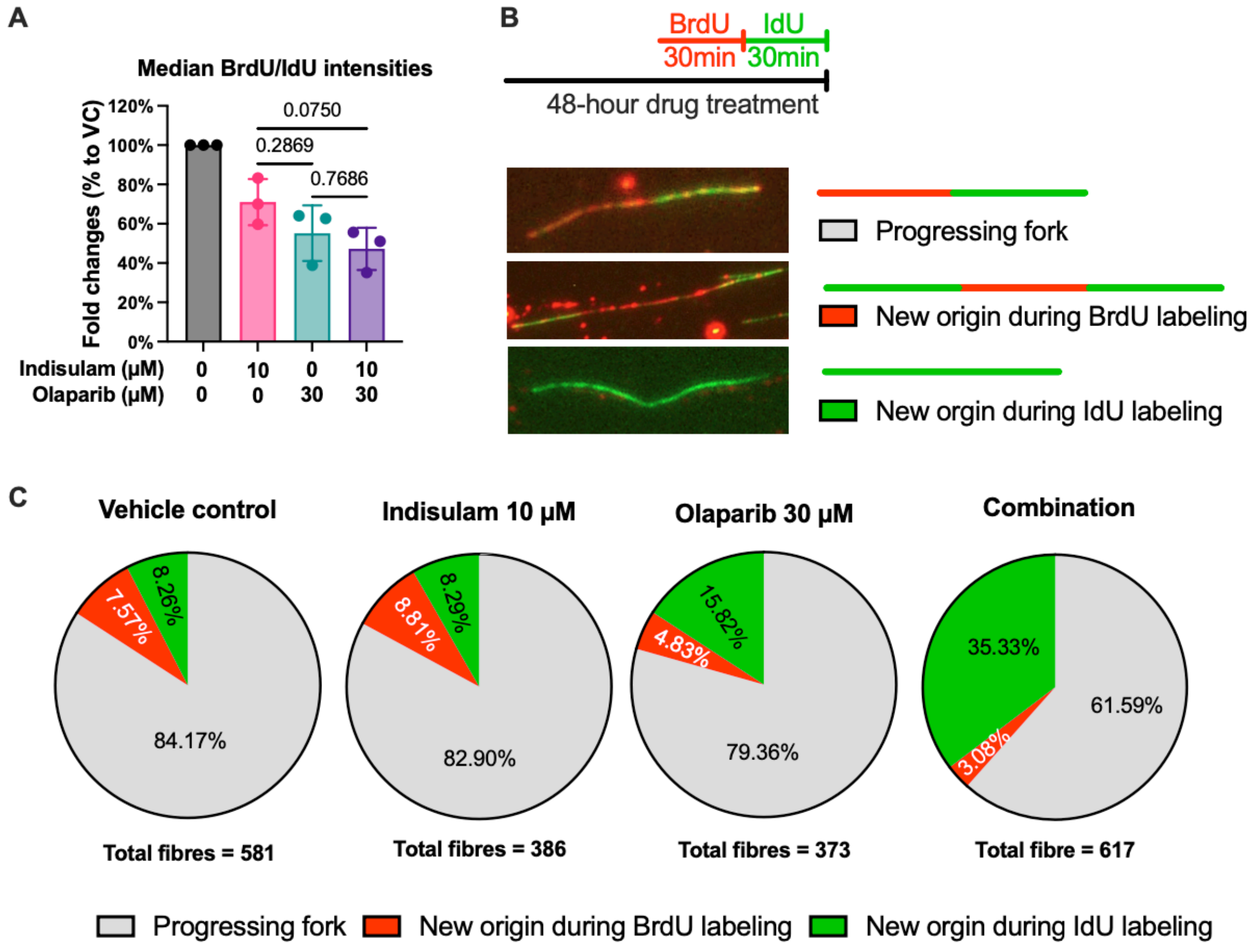


**Fig. S11. The combination of indisulam and olaparib causes DNA replication abnormalities. (A)** Quantification of median nucleotide intensities in the cell cycle analysis. Ordinary one-way ANOVA and Šídák's multiple comparisons test were used for statistical analysis, and p values are shown. Related to Fig. 5C. **(B)** The experimental design of DNA fiber analysis and examples of different types of DNA fibers. **(C)** The proportion of progression fork and new origins in KURAMOCHI cells after indicated drug treatments for 48 hours (*n*=1).

15. T. C. Chou, N. Martin. (ComboSyn, Inc., New York, USA, 2005).

16. N. Pillay, A. Tighe, L. Nelson, S. Littler, C. Coulson-Gilmer, N. Bah, A. Golder, B. Bakker, D. C. J. Spierings, D. I. James, K. M. Smith, A. M. Jordan, R. D. Morgan, D. J. Ogilvie, F. Foijer, D. A. Jackson, S. S. Taylor, DNA Replication Vulnerabilities Render Ovarian Cancer Cells Sensitive to Poly(ADP-Ribose) Glycohydrolase Inhibitors. *Cancer cell* **35**, 519-533.e518 (2019).

17. K. Cong, M. Peng, A. N. Kousholt, W. T. C. Lee, S. Lee, S. Nayak, J. Krais, P. S. VanderVere-Carozza, K. S. Pawelczak, J. Calvo, N. J. Panzarino, J. J. Turchi, N. Johnson, J. Jonkers, E. Rothenberg, S. B. Cantor, Replication gaps are a key determinant of PARP inhibitor synthetic lethality with BRCA deficiency. *Mol Cell* **81**, 3128-3144.e3127 (2021).
